# Multispecies autocatalytic RNA reaction networks in coacervates

**DOI:** 10.1101/2022.11.01.514660

**Authors:** Sandeep Ameta, Manoj Kumar, Nayan Chakraborty, Yoshiya J Matsubara, S Prashanth, Dhanush Gandavadi, Shashi Thutupalli

## Abstract

Robust and dynamic localization of self-reproducing autocatalytic chemistries is a key step in the realization of heritable and evolvable chemical systems. While autocatalytic chemical reaction networks already possess attributes such as heritable self-reproduction and evolvability, localizing functional multispecies networks within complex primitive phases, such as coacervates, has remained unexplored. Here, we show the self-reproduction of an RNA system within charge-rich coacervates where catalytic RNAs are produced by the autocatalytic assembly of constituent smaller RNA fragments. We systematically demonstrate the catalytic assembly of active ribozymes within phase-separated coacervates — both in micron sized droplets as well as a coalesced macrophase, underscoring the facility of the complex, charge-rich phase to support these reactions in multiple configurations. By constructing multispecies reaction networks, we show that these newly assembled molecules are active, participating both in self- and cross-catalysis within the coacervates. Finally, these collectively autocatalytic reaction networks endow unique compositional identities to the coacervates which in turn transiently protect the identity against external perturbations, due to differential molecular transport and reaction rates. Our results establish a compartmentalised chemical system possessing a compositional identity possessing a balance between robustness and variability required for chemical evolution.

## Introduction

Oparin and Haldane Oparin et al. (1957); Haldane (1929) hypothesized complex coacervates as self-organized chemical microreactors – spatial organizers of chemistries in a “primordial soup” – giving rise to some of the very first precursors of life. Such ‘primitive’ compartments could form a propagating (growing and dividing) core of a concentrated set of chemicals while still allowing for the exchange of material with the external environment and maintaining the system out-of-equilibrium van Harren et al. (2020). In order for such a primitive compartmentalized physico-chemical system to exhibit life-like properties, it must have a robust, heritable identity that is propagated by growth and division, allowing it to participate in Darwinian-like chemical evolution Lewontin (1970); Godfrey-Smith (2007); Adamski et al. (2020); Ameta et al. (2021a); Vasas et al. (2012); Joyce and Szostak (2018). Recently, chemically driven phase separation dynamics has been proposed to cause liquid-like coacervate droplets to grow and divide at characteristic sizes Zwicker et al. (2017), and chemical activity-driven growth of coacervate droplets has also been explored experimentally Nakashima et al. (2021). However, endowing such primitive compartments with a robust and heritable identity that has the potential for chemical evolution, remains an open challenge.

Compartmental identity can be of two broad types: (i) by encapsulating unique template molecules (coded polymers) that they carry (*e.g*. DNA in extant biological cells) or (ii) by a chemical composition – a so-called “composome” Segré et al. (2000) – comprising the compartment (*e.g*. the lipid composition of a vesicle or the relative abundance of the species of a chemical reaction). Heritable reproduction of such a compartmental entity involves either (i) replication of the template molecules coupled with the growth and division of the compartment or (ii) the generation of a second compartment with a chemical composition identical to that of the original one. While the compartment itself can be comprised of sufficiently complex molecules, it is widely argued that a templated-replicator molecular system and the maintenance of its fidelity to overcome replication errors Eigen (1971); Eigen and Schuster (1977); Dyson (1985) is contingent upon other sophisticated mechanisms (molecules), making the spontaneous emergence of such a templated-replicator – from a primitive chemical mixture – highly unlikely. It is however expected that chemical evolutionary dynamics can lead to emergence and maintenance of such a templated-replicator and associated mechanisms from simpler chemical systems Vasas et al. (2012). Such an alternate (or a predecessor) mechanism for identity establishment, replication and chemical evolution are the so-called autocatalytic chemical reaction networks (ACSs) – reaction networks in which the molecules comprising the network are collectively synthesised from a pool of chemical substrates (food) Kauffman (1986); Hordijk and Steel (2012). The steady-state chemical composition of such an ACS network – its composome – serves as a chemical identity for the network and, by extension, to any compartment that encloses the chemical network. Such autocatalytic networks have also been envisaged to emerge from random chemical pools and possess properties critical for Darwinian-like chemical evolution Nghe et al. (2015); Ameta et al. (2021a); Hordijk et al. (2012); Szathmáry and Demeter (1987); Vasas et al. (2012); Wołos et al. (2020).

Localizing ACS chemistries into dynamic compartments can open up the possibilities of a life-like evolvable chemical system — while the compartment protects the enclosed chemistry, a dynamic flux across the compartment boundary can sustain a balance between the ACS network robustness and possibility for change *i.e*. evolvability. However, the formation and preservation of ACSs requires not only the synthesis of entities capable of autocatalysis but also their interaction with similar entities to form cooperative cross-catalytic networks, all within the confines of a compartment. While ACSs have been shown to form using diverse chemistries, those formed from catalytic molecules such as RNA are of particular interest due to the possibility of them being at a crucial stage in chemical evolution Gilbert (1986), especially leading to longer and more complex information carrying polymers, and eventually to the emergence of templated replication.

While contemporary biological systems enforce our view of compartmentalization to round, spherical entities, in general, abiotic spatialization can occur in the form of eutectic phases Menor-Salván and Marín-Yaseli (2012), pores in hydrothermal vents Martin et al. (2008); Deamer and Georgiou (2015), mineral surfaces Hazen and Sverjensky (2010), vesicles Chen and Walde (2010), phase-separated compartments Martin (2019) and even hydrodynamic flow structures Krieger et al. (2020). Phase-separated compartments provide a versatile and diverse setting for chemical dynamics Ianeselli et al. (2022) – for example, they can comprise of liquid-like droplets, gel-like particles, or even condensed macrophases resulting from the coalescence of these structures Ianeselli et al. (2022) (Fig. 1A). In order to establish coacervate-based compartments as evolvable entities, it is important to demonstrate self-reproducing chemistries in multiple phase-separated (coacervate) scenarios.

**Figure 1.**
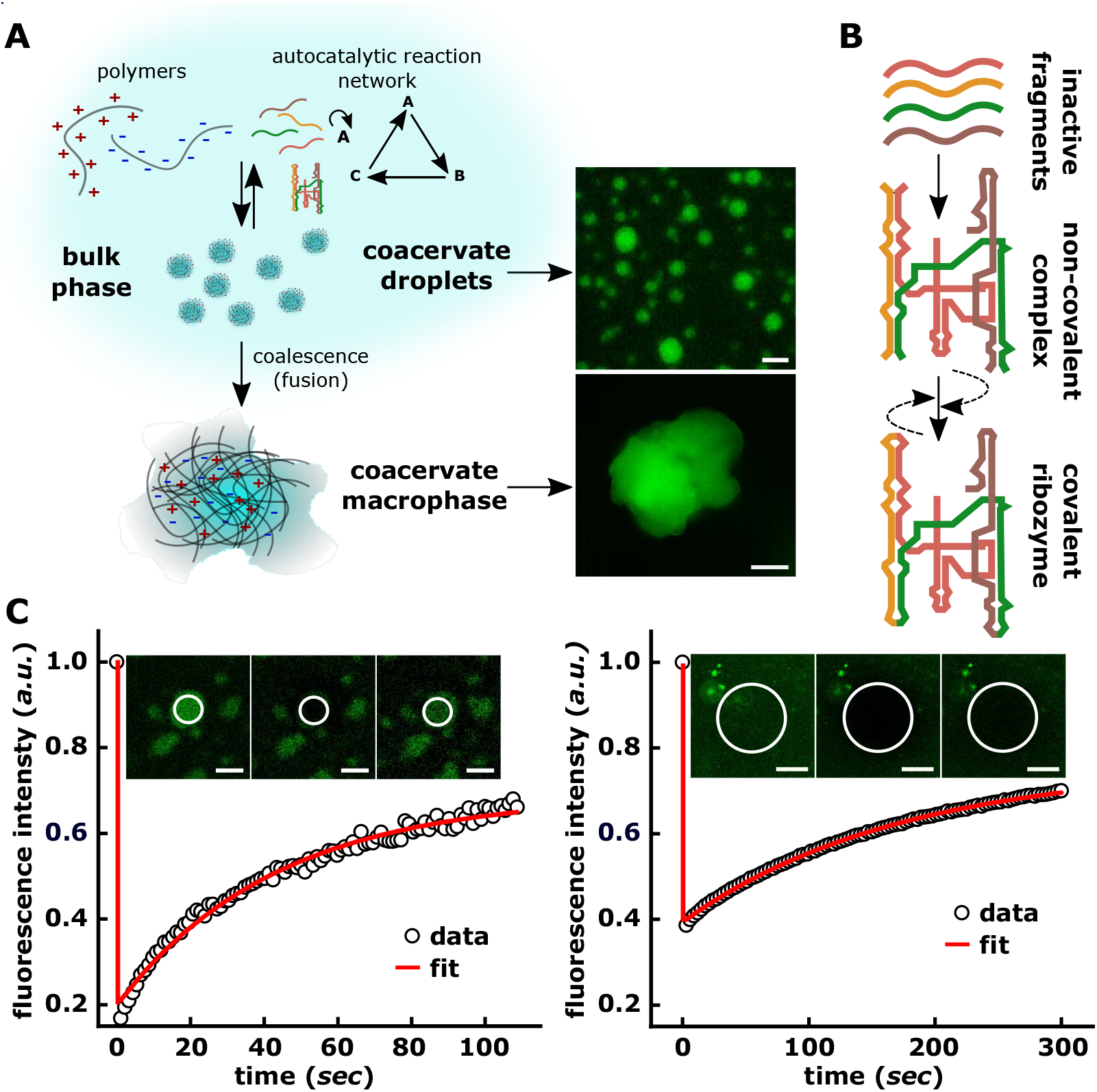
Conceptual schematic of our work involving coacervate compartments and a RNA ribozyme system. **A**. Schematic representation of phase-separated compartments formed by interaction between oppositely charged polymers (PAA and spermine, see Materials and Methods). These phase-separated compartments can exist as spatially isolated, micron sized ‘coacervate droplets’ in solution or can coalesce into a single consolidated ‘coacervate macrophase’. Both these coacervate environments support autocatalytic RNA self-assembly and reaction network formation. Microscopy images (green fluorescence channel) of PAA-spermine coacervate droplets (*top right*, scale bar 5 *μ*m) and coacervate macrophase (*bottom right*, scale bar 10 *μm*). The fluorescence is due to 30nt long oligonucleotides labeled with Alexa-488 added during the coacervation step (see Materials and Methods, Supplementary Table 1). **B**. Schematic showing the self-assembly of *Azoarcus* covalent ribozyme (WXYZ, ~200nt) from its inactive substrate RNA fragments (*red* : W ~65nt, *yellow* : X ~43nt, *blue*: Y ~52nt, *brown*: Z ~55nt) Hayden and Lehman (2006); Hayden et al. (2008). The 5’-end of W and 3’-end of W, X, and Y contains 3nt long recognition elements annotated as “IGS” and “tag”, respectively. When mixed together, the substrate RNA fragments rapidly self-assemble to form a non-covalent complex (catalytically active) which is converted to a covalent ribozyme (catalytically active, WXYZ). The dashed arrow denotes catalytic feedback from the non-covalent as well as covalent catalysts. **C**. FRAP (fluorescence recovery after photo-bleaching) is used to characterise the **C**. coacervate droplets and **D**. coacervare macrophase. Alexa-488 labeled 30nt long oligonucleotide inside the coacervate droplets are used to perform the experiment at 48°C. Insets show the sample before and after the photo-bleaching, scale bar 5 *μm*.

However, achieving functional RNA catalytic assembly and reaction networks within the charge-rich environments of complex coacervates is a challenging task. Some of the foundational work involves demonstrations of RNA exchange between a coacervate and its environment Jia et al. (2014) and the catalytic activity of RNA cleavage and template-based polymerization in coacervates or ATPS (aqueous two-phase systems) Drobot et al. (2018); Poudyal et al. (2019); Strulson et al. (2012). Furthermore, several recent studies report some critical steps towards the construction of an evolvable coacervate-based protocell, *e.g*., catalytic assembly of longer RNA in condensates Le Vay et al. (2021), RNA reproduction via ligase activity Iglesias-Artola et al. (2022); Salditt et al.; Salibi et al.. However, the (auto-)catalytic assembly of longer (larger) molecules from constituent fragments, the creation of reaction networks and achieving self-reproduction remain explored.

Here, we perform experiments with fragments of the *Azoarcus* group I intron ribozyme Reinhold-Hurek and Shub (1992) to demonstrate autocatalytic assembly of longer catalytic RNA molecules within complex, charge-rich coacervate phases (Fig: 1B). (i) We demonstrate the self-assembly of large RNA catalytic molecules from constituent smaller fragments inside a charge-rich coacervate macrophase as well as spatially isolated droplets; (ii) this assembly can be achieved in both a self-autocatalytic and collective cross-catalytic fashion, allowing us to form ACSs within coacervates, (iii) these ACS reaction networks confer a chemical compositional identity to the coacervate compartments, and (iv) despite the lack of an impermeable boundary, both of the dynamic compartments — coacervate droplets and macrophase — transiently protect the enclosed reaction networks (and by extension, the chemical identity) from external perturbation. Altogether, by combining RNA autocatalytic networks with dynamic phase-separated coacervate compartments, our work opens the possibilities for creating primitive chemical units, that can participate in self-reproduction, growth and division Zwicker et al. (2017); Nakashima et al. (2021); Ianeselli et al. (2022) and adaptive evolution Ameta et al. (2021).

## Results

The *Azoarcus* ribozyme, WXYZ, is a ~200-nucleotide (nt) RNA (Fig. 1B) which can be split into inactive RNA fragments of varying lengths (two fragments Yeates et al. (2016, 2017); Ameta et al. (2021), four fragments Hayden and Lehman (2006), five fragments Jayathilaka and Lehman (2018)). These fragments can covalently assemble in full-length active ribozymes in the presence of Mg^2+^ ions using both catabolic Arsène et al. (2018) and anabolic Hayden et al. (2008) approaches via recombination reactions Hayden and Lehman (2006). The assembly occurs in an autocatalytic manner *i.e*. the self-assembled ribozymes catalyze the subsequent assembly of fragments into other functional ribozymes to form autocatalytic networks capable of collective reproduction Hayden et al. (2008); Vaidya et al. (2012). Further, the assembly is guided by specific base-pair interactions between a 3nt long internal guide sequence (IGS, “GUG” in wild-type group I intron) at the 5’-end of the ribozyme with the complementary 3nt stretch (tag “CAU”) at the 3’ -end of the substrate RNA fragments. Variations in the IGS and tag sequences allow for the creation of a diversity of ribozymes that can be used to build cross-catalytic reaction networks Vaidya et al. (2012); Ameta et al. (2021).

We prepared coacervates using a negatively-charged long chain poly-acrylic acid (PAA) and positively-charged spermine creating liquid-like droplets of varying size (0.5-5 *μ*m) (Fig. 1A, Materials and Methods, and Supplementary Fig. S1). The coacervate phase is a charge-rich environment, with complex rheological properties that can not only affect the mobility of molecules into (and within) the condensed phase but also their reaction dynamics. Therefore, at first, by using fluorescently labelled oligonucleotides (Fig. 1A, *right*), we confirmed that negatively charged polymers (such as RNA or DNA) can spontaneously partition into the coacervate phase. These samples are then used to characterize the dynamics within the compartment – FRAP (fluorescence recovery after photobleaching) measurements on the PAA-spermine coacervate droplet as well as macrophase confirm diffusive dynamics of the oligonucleotides and dynamic exchange with the external environment (*droplet* : recovery time *τ* = 21.3 ± 3.0 s; diffusivity *D* = 0.1 ± 0.02 *μ*m^2^s^−1^, *macrophase*: recovery time *τ* = 172.5 ± 4.0 s and diffusivity *D* = 98 ± 1.8 * 10^−3^*μ*m^2^s^−1^, Fig. 1C, D; Materials and Methods).

The partitioning of RNA molecules into the coacervates does not however guarantee their functionality, *i.e*., catalytic activity in the charge-rich and confining environment of coacervate droplets as well as that of the coalesced macrophase Ianeselli et al. (2022). Identifying and quantifying any ribozyme activity inside such a charge-rich environment necessitates the careful distinction between reactions occurring inside the coacervate droplets, macrophase, and in the bulk solution phase (Fig. 1A). We, therefore, developed a protocol to separate the bulk of the solution from the coacervate phase and systematically analysed the ribozyme activity within droplets as well as macrophase (Materials and Methods and Supplementary Fig. S2). Briefly, coacervates droplets were prepared encapsulating all the four RNA fragments (W, X, Y, and Z; Fig. 2A) and incubated at 48°C. Indeed droplets are stable at such higher temperatures of the reaction (Supplementary Fig. S3). To measure activity from coacervate droplets, the bulk aqueous phase was removed using brief centrifugation after the incubation (Materials and Methods; Supplementary Fig. S2) and separated phase was analyzed using polyacrylamide gels (Materials and Methods). Similarly for the coacervate macrophase, the bulk aqueous phase was separated prior to the incubation at 48°C and only the separated coacervate macrophase (with no bulk solution around) was incubated ensuring RNA catalysis inside the coacervate macrophase.

**Figure 2.**
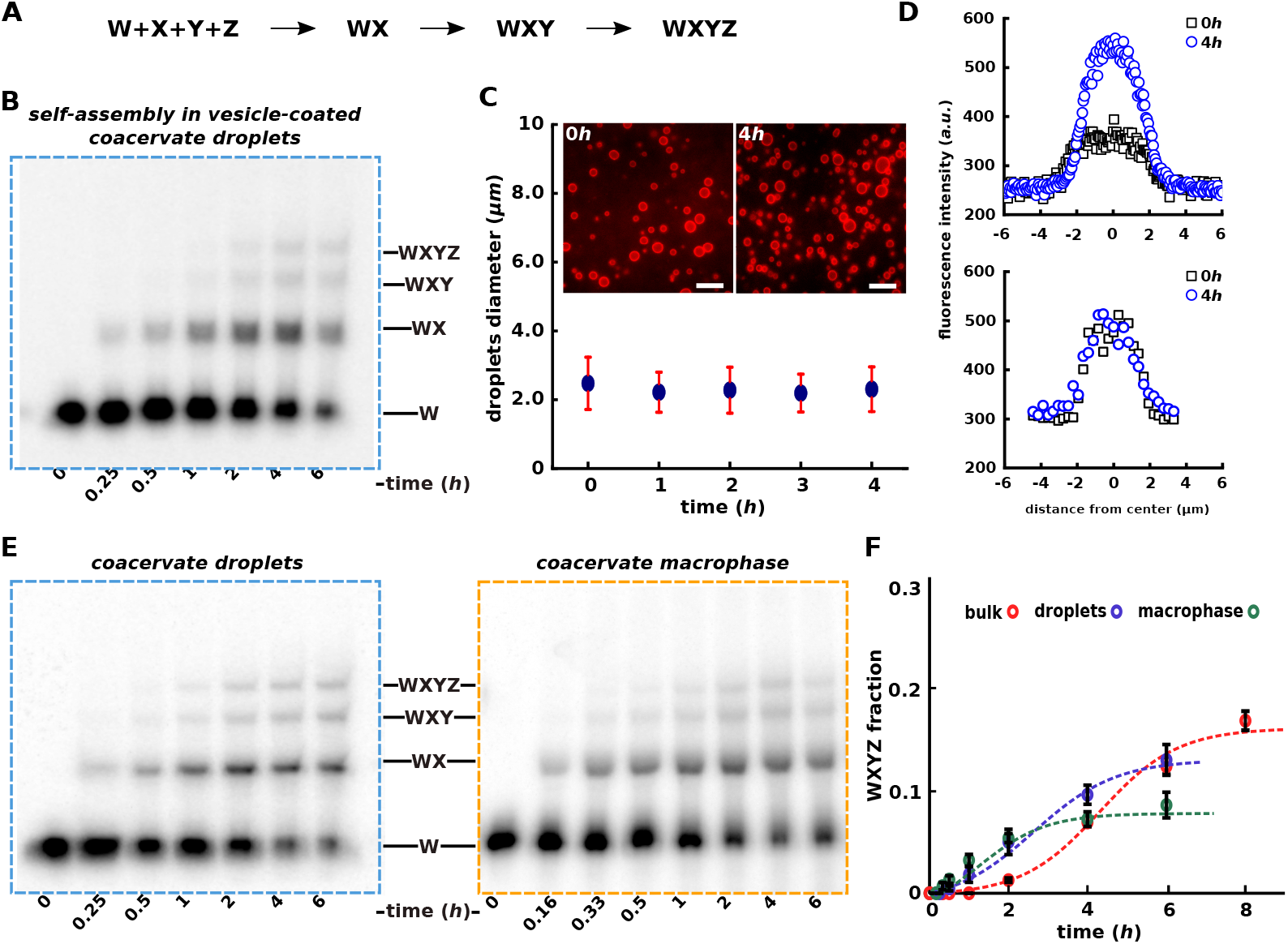
Autocatalytic self-assembly of the *Azoarcus* ribozyme in coacervates. **A**. Schematic showing the systematic assembly of covalent ribozyme (WXYZ) from smaller inactive RNA fragments W, X, Y, Z via WX, WXY intermediates. **B**. Polyacrylamide gel showing the self-assembly of *Azoarcus* ribozyme (WXYZ) from its small RNA fragments (W, X, Y, Z) inside the vesicle-coated coacervate droplets demonstrating that WXYZ formation indeed occurs inside the micron-size droplets. Here coacervate droplets were prepared together with RNA fragments (W, X, Y, and Z) and then coated with lipid-vesicles (DOPC) prior to incubating at 48°C and analyzed over polyacrylamide gel (see Materials and Methods). **C**. Graph showing the stability of vesicle-coated coacervates during the reaction time-course shown in **B**. Here average droplet diameter is plotted over the time. The microscopy images of vesicle-coated coacervate droplets (at 0*h* and 4*h*) are shown in inset. The fluorescence is due to the doping of 1,2-dioleoyl-sn-glycero-3-phosphoethanolamine-N-(lissamine rhodamine B sulfonyl) with the DOPC lipid (see Materials and Methods). **D**. Graph showing the leakiness of the vesicle-coated coacervate used for the self-assembly reaction shown in **B**. Here leakiness is tested as increase in fluorescence intensity of the coacervate droplets due to the diffusion of a 30nt Alexa-488 DNA oligonucleotide (same as used in FRAP studies above, Fig. 1) from bulk solution to the vesicle-coacervate droplets. Leakiness is measured either for the 30nt Alexa-488 DNA oligonucleotide alone (*top*) or hybridized to the ~200nt *Azoarcus* ribozyme (*bottom*). See Materials and Methods for the details. **E**. Polyacrylamide gels showing the time course for the formation of full-length product (covalent ribozyme, WXYZ) starting from smaller RNA fragments inside coacervate droplets (*left*) as well as inside coacervate macrophase (*right*). These reactions are carried out in the presence of a (*γ*^32^*P*) labelled W RNA fragment and analyzed via 12% denaturing PAGE (see Materials and Methods). **F**. Time-courses showing the autocatalytic self-assembly of WXYZ ribozyme from W, X, Y, and Z RNA fragments in coacervate macrophase (green circles), coacervate droplets (blue circles) and as well as in bulk aqueous phase control (red circles). Reactions are done by encapsulating the substrate fragments in coacervates, separating the macrophase, or by incubating the droplets directly at 48°C (Supplementary Fig. S2 and Materials and Methods). All the time-courses are measured in triplicates and mean WXYZ product formation is plotted along with standard error. The time-course data is also analyzed by a kinetic model described for the *Azoarcus* assembly earlier Hayden et al. (2008) and the fitted curves are shown as dotted lines (green, blue, and red for the macrophase, droplets, and bulk aqueous phase, respectively). The details of the kinetic model are described in Materials and Methods section.

First, in order to confirm that the RNA ribozyme assembly can occur inside micron-sized coacervate droplets and to rule out the diffusion of the assembled product from bulk aqueous solution present around into the droplet, we devised an experiment in which the coacervate droplets containing RNA fragments were coated with lipid-vesicles. These lipid vesicles coating the droplets restrict the RNA molecular transport from the bulk to the droplets (Materials and Methods). Therefore, the formation of full-length WXYZ within these vesicle-coated coacervate droplets unambiguously confirms that the reactions occur inside the micron sized coacervate droplets (Fig. 2B). Additionally, we have verified that these vesicle-coated coacervate droplets are stable against coalescence (Fig. 2C) and that the longer WXYZ RNAs cannot partition from the bulk (Fig. 2D) at 48°C for longer duration. Altogether, these results affirm that the coacervate phase, even in the form of micron sized droplets, can sustain the RNA self-assembly reactions.

Next, we systematically analyzed the assembly of WXYZ ribozyme inside membraneless coacervate droplets *i.e*. without any vesicle coating as well as inside the macrophase. Following the same protocol mentioned above, in both the cases, appearance of slower-moving products on a polyacrylamide gel corresponding to the size of full-length WXYZ, confirms the assembly of longer RNAs (*Azoarcus* ribozyme, Fig. 2E). The autocatalytic nature of the fragment assembly is evident in all cases: bulk solution phase, coacervate droplets as well as coacervate macrophase, as the time course of the WXYZ product formation follows a sigmoidal trend that is expected for autocatalytic growth in a closed system Hayden and Lehman (2006) (Fig. 2F, red, blue, and green data-points, see Supplementary Text 1). Additionally, we have also analyzed the four-fragment assembly of WXYZ in presence of product of the reaction (full-length covalent WXYZ) seeded at the start of the reaction confirming the positive feedback in the assembly (Supplementary Fig. S4).

In order to explain the kinetic behaviour of the assembly, we used a kinetic model, simplified from one described for the *Azoarcus* assembly previously Hayden et al. (2008), to analyse the time-course data (Fig. 2F, red, blue and green dotted lines). Briefly, we considered the instantaneous assembly of a non-covalent complex from the four fragments (W, X, Y and Z) Vaidya et al. (2012); Ameta et al. (2021), and a (reversible) conversion reaction of W:X:Y:Z into a covalent catalyst (WXYZ) catalyzed by both WXYZ and W:X:Y:Z with different catalytic efficiencies, *k_a_* and *k_b_*, respectively (details in the Materials and Methods section). The model parametrizes only three quantities: the strength of positive feedback due to the formation of a catalytic product *a*, the rate of the background reaction (catalyzed by the non-covalent complexes) *b*, and the product fraction at equilibrium *c* (Supplementary Table 2). The data from all the different experiments fit equally well using this model and a sigmoidal nature of the time course is evident (Fig. 2F and Supplementary Fig. S5), thus unambiguously demonstrating the autocatalytic nature of the RNA ribozyme assembly. In addition to this kinetic modeling of the time course data, by measuring the initial reaction rates of the WXYZ assembly from two fragments (WXY and Z) seeded with different initial concentrations of WXYZ as described earlier Yeates et al. (2016); Ameta et al. (2021) (Supplementary Fig. S6), we confirmed positive feedback from the product of the reaction (WXYZ), further verifying the autocatalytic nature of the WXYZ assembly (Supplementary Text 1).

Altogether, these results show that coacervate compartments support the autocatalytic assembly of catalytic RNAs from smaller RNA fragments despite the condensed and charge-rich conditions — it must be emphasized here that this assembly not only requires the coming together of the various fragments but also the final assembled structure to adopt a functional secondary structure for further catalytic action.

Another notable feature of the reactions inside the coacervate compartments is the rate enhancement of the assembly, as also observed earlier for other catalytic RNAs Poudyal et al. (2019). When compared with the same self-assembly reaction in the absence of coacervation (bulk control), the formation of the assembled ribozyme product is significantly faster in coacervate compartments, evident by a shorter ‘lag-phase’ (Fig. 2F). Reactions within the coacervate macrophase are faster than those in coacervate droplets — this is to be expected since the surface area to volume ratio of the droplets is much higher than that for the macrophase which prevents any dynamic exchange with the bulk that can slow the effective rates of the reactions. These reaction rate enhancements are evident in the kinetic parameters (Supplementary Table 2) obtained from fits of the model to the data: while the rates of the background reaction *k_b_* in the coacervate compartments are significantly higher than those in the bulk phase, the reaction rate catalyzed by the covalent catalyst *k_a_* does not indicate such an enhancement. A similar enhancement is observed in the initial production rate measured in WXYZ assembly from two fragments (Supplementary Fig. S6). Spermine, which is present abundantly in the coacervate phase is known to stabilize RNA structures Szer (1966), could lead to rate enhancements that observed here; controls where reactions were carried out with and without spermine, however, rule out that possibility as similar amount of product formed in both the cases (Supplementary Fig. S7). Instead, we find that the increase in the reaction rates is attributable to the increase of the RNA concentration within the coacervate phase — indeed, we observe a ~ 50 fold increase in the RNA amount inside the coacervate macrophase phase compared to that in the aqueous phase alone (Supplementary Fig. S8).

We next turn our attention to the formation of cross-catalytic networks of ribozymes inside the coacervate compartments. Cross-catalysis, as discussed earlier, is important for the construction of larger and more diverse autocatalytic reaction networks with properties crucial for network robustness, self-reproduction, and Darwinian-like evolution of autocatalytic chemical systems Ameta et al. (2021,a). In addition, Darwinian evolution also requires network formation inside compartments as it allows for competition between compartments containing distinct networks. The most striking manifestation of cross-catalysis involves a poor self-assembling species, the formation of which can be catalysed by other chemical species — the growth of such a species is significantly enhanced due to cross-catalysis and is slow otherwise.

To test this, we chose a poor self-assembler (IGS-tag combination of GUG and CUU respectively; denoted as UU species Yeates et al. (2016)) compared for example with another good self-assembling species (for example, UA; Supplementary Fig. S9). We then designed cross-catalytic networks of different architectures and sizes (containing either two, three or four nodes) in such a way that the synthesis of the poor self-assembler UU is dependent on the catalysis by other members of the network (Fig. 3). For example, in the case of two nodes network, both UU and AA are poor self-assembler but can cross-catalyze each other by U→A link and A→U link, respectively. To this end, all the RNA fragments required for a particular catalytic reaction network are encapsulated in the coacervate and analyzed in droplets as well as macrophase. A radioactive labeled WXY fragment for the formation of the UU species is used to monitor the growth of the node corresponding to the poor self-assembling species in each of the networks. The time-courses, denoting the assembly of the UU species, clearly indicate significant enhancement in both coacervate droplets and macrophase (Fig. 3A, B). In the case of coacervate macrophase up to 50% of the substrate converted into the product when UU is connected in network compared to 5% when UU is alone ((Fig. 3B). These results show the assembly when the poorly self-assembling species is part of a cross-catalytic network; this is a clear demonstration of cross-catalytic reaction networks within the coacervate droplet and macrophase. Further, the production of the UU species due to the catalysis of two catalytic links (for *e.g*., in a four-node network) is enhanced when compared to production due to one catalytic link alone (for *e.g*., in two-, three-node networks); this is further confirmation of cross-catalytic network formation (Fig. 3B, compare blue with orange and red curves).

**Figure 3.**
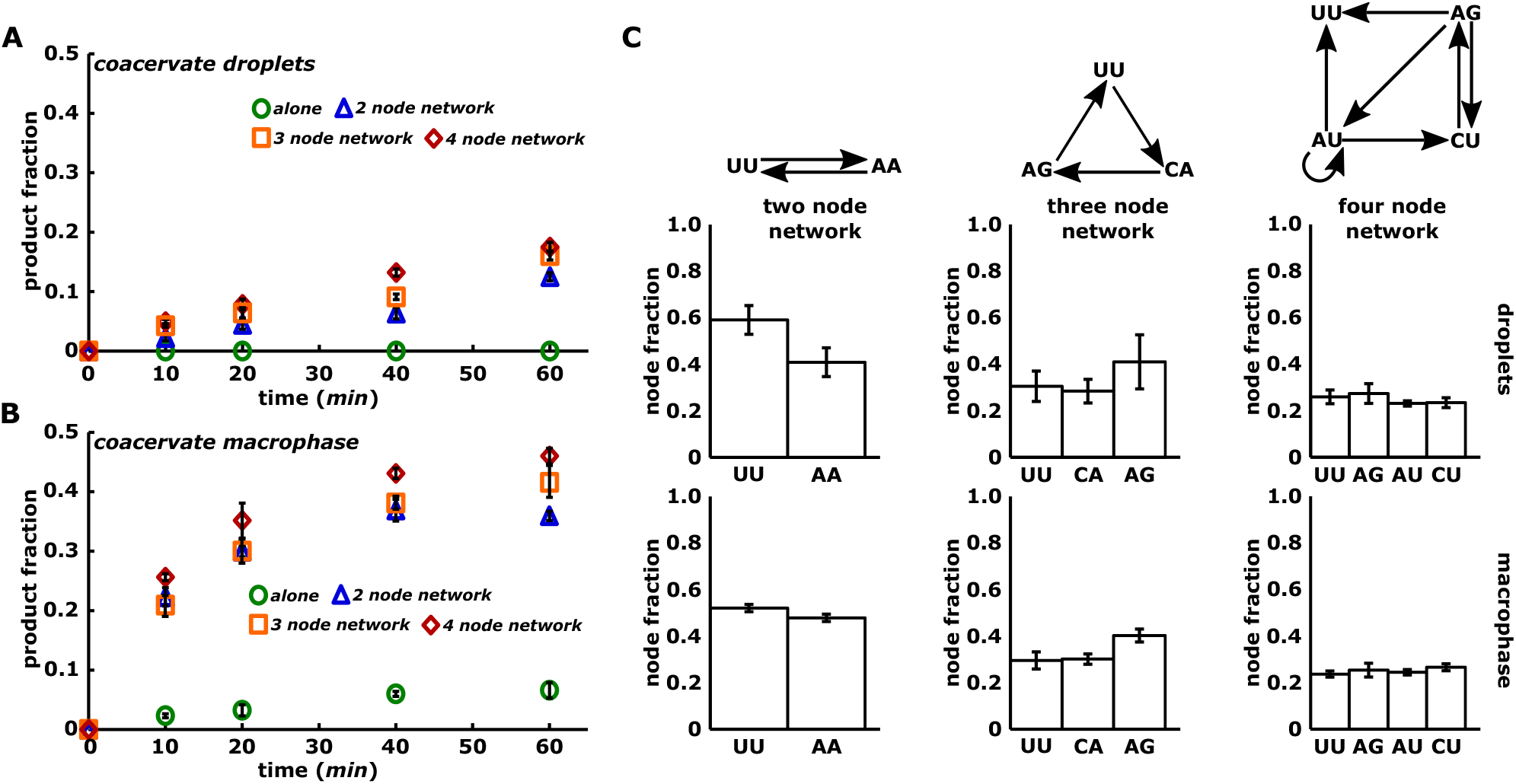
Formation of cross-catalytic networks inside coacervates. **A** Time-courses showing the assembly of node UU as single node (alone, green circle), connected in a two nodes (blue triangle), three nodes (orange square) and four nodes network (red rhombus) in coacervate droplets. Network structures are shown in **C** (*top*). **B**. Same as **A** but in the coacervate macrophase. All the time-courses are measured in triplicates and mean WXYZ product formation is plotted along with standard deviation. **C**. Bar-graphs showing the composition of two (*left column*), three (*middle column*), and four nodes network (*right column*. Network structures are shown at the top of each column. In these networks, arrow represents a directed edge showing the catalysis of a downstream node (WXYZ catalyst) by upstream node(s) (WXYZ catalyst) from its respective substrate fragments Vaidya et al. (2012); Yeates et al. (2017); Ameta et al. (2021). Here, compositions are represented as relative fraction of each network species (node) in the network and measured by quantifying the amount of each species formed in the network. All measurements were performed in triplicates and mean WXYZ product formation of each is plotted along with standard deviation.

Autocatalytic chemical reaction networks within the coacervate compartments are a natural way to the establishment of a compositional identity for the compartments Ameta et al. (2021,a). Therefore, for two-, three- and four-node reaction networks (Fig. 3C), compositions were measured by encapsulating all the substrate fragments required for a network and quantifying the formation of all the nodes individually (Materials and Methods). The measurements reveal that the network composition (relative fraction of network species) (Fig. 3C) are maintained within the coacervate droplets and as well as in coacervate macrophase, despite the charge-rich environment, and the increased RNA concentration within the compartment; the relative proportions of the nodes in networks are indeed well correlated with those measured in the aqueous phase alone (Supplementary Fig. S10). Slight differences in some of the compositions may be due to the concentration dependant nonlinearities of the reaction network kinetics and the transport within the condensed compartments.

Chemical reaction networks provide the potential for growth and adaptation dynamics for the compartment in which they are encapsulated. Further, compartmentalization can be crucial for the subtle balance between the protection of the chemical properties of the compartment and the external chemical perturbations that can drive chemical evolution. Such a balance between robustness of the compositional identity and ability to explore new chemical states have been shown to be crucial for Darwinian evolution to occur in the chemical networks Ameta et al. (2021). Thus, we sought if such a balance is possible in our system *i.e*. providing a robustness to the chemical identity against parasitic perturbations while still maintaining chemical dynamicity (that allows for the networks to expand by the addition of new species/nodes). Perturbation scenarios can be easily constructed using phase-separated compartments since they can be maintained as individual entities and as well as, are amenable to dynamic fusion-fission events that allow for the mixing of information, that can aid in sustaining evolving chemical systems Ianeselli et al. (2022). However, despite such fusion events we wondered whether membrane-less compartment system such as coacervates can impede the perturbation to the chemical composition of autocatalytic reaction networks.

To test this, we designed a two node reaction network containing the UA and UG ribozyme species (Supplementary Text 2). The UA and UG WXYZ ribozymes are connected by single catalytic edge from UG to UA via the U→A link Yeates et al. (2016). WXYZ UG ribozyme catalyses the synthesis of WXYZ UA which in addition, is catalyzed by a self-loop (U→A link, Fig. 4A, *top left*). UG is also self-catalyzed by a weak wobble base-pair link (U→G link, not shown here). As expected, when network species were measured in the bulk aqueous phase, this arrangement resulted in a composition with a higher proportion of the UA species compared to UG species (Fig. 4A, *bottom left*). We then ‘perturbed’ the network by the addition of a catalytic species CA that catalyses the UG node (with the C→G link, Fig. 4A, *top right*). CA perturbs the target node strongly since it catalyses an isolated species, which is only weakly catalyzed, with a strong catalytic link (C→G link) Ameta et al. (2021). The perturbation is indeed evident from the change in chemical composition of the network in the bulk aqueous phase — the compositional ranking of the UG and UA species is flipped upon the addition of the CA species which catalyzes the synthesis of UG species (Fig. 4A, *bottom right*; compare the composition in *bottom left* with *bottom right*). While such a disturbance of the composition is expected in a well-mixed solution, to test this in the coacervate compartments we devised two situations: (a) the network is encapsulated in the coacervate droplet (Fig. 4B), and (b) networks inside the coacervate macrophase (Fig. 4C). When the network was encapsulated inside coacervate droplets and CA encapsulated droplets were used for perturbation, we observed the network composition is protected, i.e., the compartment was indeed able to render protection to the network composition as the added perturbation does not result in a change in the compositional identity (Fig. 4B). Similarly, for the macrophase, perturbation was achieved by adding solution CA species on top of the separated coacervate macrophase and then incubated. Here too, the coacervate macrophase compartment rendered protection to the compositional identity (Fig. 4C). Unlike in surfactant-stabilized aqueous droplets in an oil phase Ameta et al. (2021), or vesicles formed by lipids, which are impermeable to large RNA fragments altogether, the network robustness that is conferred by the coacervates is transient — the compartments arise due to a liquid-liquid phase-separation and lack an impermeable boundary. Indeed, upon a longer incubation (4 h compared to 30 min used above) of the system under perturbation, the chemical composition within the coacervate is modified (Supplementary Fig. S11). Perturbation time-scales in such condensed coacervate system would depend on multiple parameters like length of polymers involved, net charge, structure of the nucleic acids and would require detailed exploration. To be a good model protocell, a coacervate compartment must not only have a robust self-replicating chemistry inside it, but also be able to transport molecules into and out of the compartment. Therefore any associated perturbation to the network robustness can happen on timescales that are long enough for selection to act — one of the possible ways is implementing non-equilibrium scenarios with dynamic fusion and fission of droplets as was recently reported Ianeselli et al. (2022). Altogether, our experiments do indeed show a clear separation of reaction timescales (network identities get perturbed over the course of hours) and diffusion timescales (happening on the timescale of a few minutes as measured from the molecular diffusion), which is crucial for a system to exhibit both heritability and evolvability.

**Figure 4.**
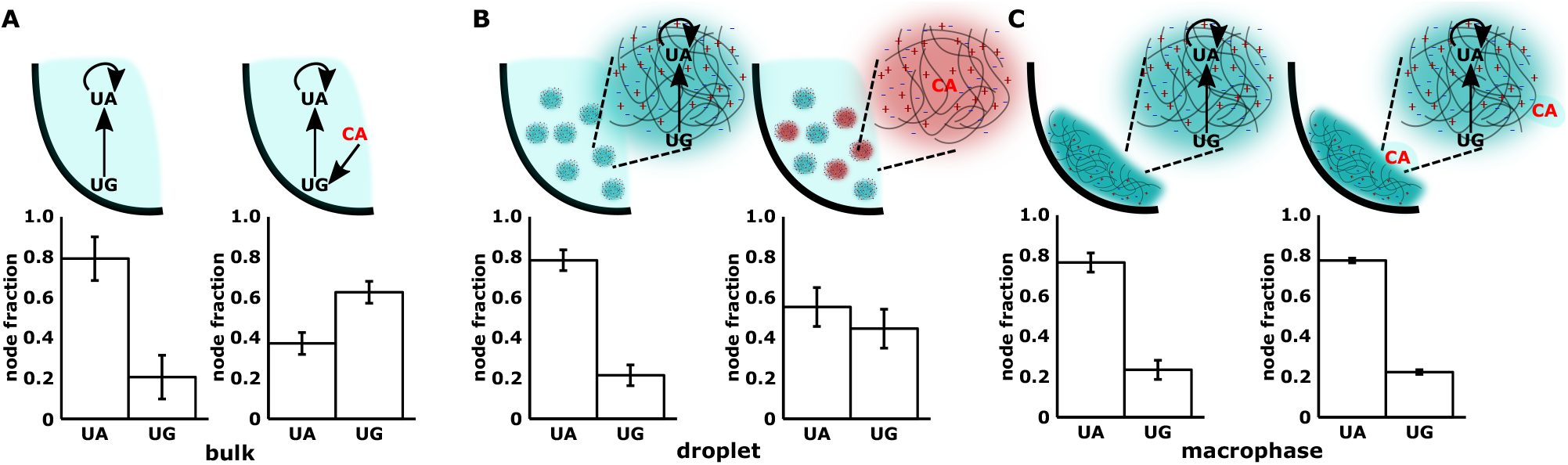
Robustness of cross-catalytic RNA networks in coacervates. Compositions are measured for a two node cross-catalytic network in the absence or presence of a perturbing species under three different conditions. In this network, UA is self-catalyzed, as well as catalyzed by UG (via U→A link, *panel* A, *top left*). UG is also poorly self-catalyzed via U→G (*not shown*). The perturbation is caused by adding a strong node (CA, shown in red) which catalyzes the synthesis of UG node (via C→G link, *panel* A, *top right*. **A**. *bulk* : Network is formed in the absence of any polymer and compositions are measured without perturbation (*bottom left*) as well as with perturbation (CA, *bottom right*). **B**. Same as **A** but in coacervate droplets. Here, perturbing species (CA) is also encapsulated inside the coacervate droplets followed by mixing with droplets carrying the two-node network in 1:1 ratio, *left* without perturbation, *right* with perturbation. **C**. Same as **B** but in coacervate macrophase. The perturbation is carried out by adding 0.5 *μM* solution of CA on top of macrophase, *left* without perturbation, *right* with perturbation. For each condition, samples were incubated for 30 *min* and then analyzed via polyacrylamide gel electrophoresis to measure the network compositions with or without perturbation. All the measurements are done in triplicates and mean WXYZ product formation is plotted as fraction w.r.t. each node along with standard deviation.

## Discussion and Conclusions

Integrating RNA autocatalytic reaction networks with coacervate compartments, brings together three important conceptual frameworks related to the emergence of a protocell from a ‘primitive’ chemical system: (i) the RNA world hypothesis Woese (1967); Crick (1968); Gilbert (1986) or more broadly, the dynamics of sufficiently long, information carrying catalytic molecules capable of participating in complex reactions and evolving in complexity, (ii) autocatalytic reaction networks, which possess all the attributes required of a simple chemical system to display the properties of robustness, heritability and evolvability Kauffman (1986); Hordijk and Steel (2012); Ameta et al. (2021a) and (iii) dynamic compartmentalization via coacervation Oparin et al. (1957); Haldane (1929) which serves to spatio-temporally localize chemistries. The RNA world has long been suggested as a plausible stage of chemical evolution in which self-replicating RNA molecules (or in general, sufficiently complex catalytic polymers) proliferate, and evolve to give rise to more complex molecules with diverse functionalities, before the appearance and evolution of other complex life machinery such as DNA and proteins Gilbert (1986). However, the steps from the appearance of nucleotides Powner et al. (2009) to the development of complex, functional polymers such as an RNA replicase have not been fully elucidated; the emergence of RNA replicases and templated replication, for instance, is plagued by barriers such as parasites and replication errors Eigen (1971). Autocatalytic reaction networks Kauffman (1986); Ameta et al. (2021a) and compartmentalization Matsumura et al. (2016); Vasas et al. (2012) have both been proposed to ameliorate such barriers (among others Nghe et al. (2015); Ameta et al. (2021a)), to give rise to increasingly complex, functional molecules and mechanisms crucial for the emergence of a protocell. In bringing these various conceptual frameworks together in a single physical realisation *i.e*. coacervate-based compartments with autocatalytic RNA chemistries, our work not only fills a crucial gap of coupling but immediately opens up the possibilities to explore diverse chemical dynamics and evolutionary scenarios: (i) The assembly of catalytic RNAs from smaller inactive RNA fragments in such a charge-rich and condensed physical environment highlights the possibility that the nucleic acid-based self-reproducing systems can achieve fairly complicated functions in seemingly unfavorable conditions. (ii) Cross-catalysis and network formation, not only involves the self-assembly of catalytic molecules but that they can catalyze the synthesis of other network species. (ii) Notably, the enclosed autocatalytic reaction networks assembling RNA ribozymes endow the coacervate compartments with an identity which are protected from parasitic perturbations. Such transient protection afforded by the coacervate compartments to these complex ACSs, when coupled with growth and division mechanisms of the compartment Zwicker et al. (2017); Nakashima et al. (2019); Ianeselli et al. (2022), poises the whole system at a balance between robustness and flexibility, resulting in a heritable, self-reproducing unit, which might itself be considered a minimal protocell. Such a protocell naturally lays the ground for evolutionary mechanisms resembling canonical Darwinian evolution. When compared to vesicles, coacervate compartments are ‘leaky’ due to the lack of a defined boundary and hence using them in any prebiotic dynamical scenario would require that information propagation (*e.g*. via molecular diffusion) occurs faster than the percolation (replication) of the genetic material they carry. The implementation of hybrid systems based on lipid-coated coacervates, as we have also explored here, could overcome such percolation barriers Cakmak et al. (2019).

## Materials and Methods

### A. Materials

All the experiments used DNAse/RNase free water (Thermo Fisher Scientific Product No.: 10977015). Chemicals were purchased from SRL chemicals (Sisco Research Laboratories (SRL) Pvt. Ltd., India) and Sigma-Aldrich unless specified otherwise. 4-(2-Hydroxyethyl)-1-piperazinepropane-sulfonic acid (EPPS) was purchased from Alfa Aesar (Product no.: J60511). Two different type of poly acrylic acids were used (PAA): high molecular weight HMWPAA (~ >4,000,000 Da, Sigma Aldrich, Product no.: 306231) for preparing coacervates and low molecular weight LMW-PAA (~ 1,800 Da, Sigma Aldrich, Product no.: 323667) for dissolving the coacervates to prepare the samples for gel electrophoresis. Spermine with >97% purity from Sigma (Sigma Aldrich Product No.: S3256) was used for coacervation. 12% denaturing polyacrylamide gels were prepared using acrylamide (SRL chemicals, Product no.: 15657), bis-acrylamide (Sigma Aldrich, Product no.: 146072) in 19:1 ratio and 8 *M* Urea (Qualigens, Product no.: Q15985) and polymerized using TEMED (tetramethylethylenediamine, SRL chemicals, Product no.: 52145) and ammonium persulfate (SRL chemicals, Product no.: 65553). 2% agarose gels were prepared from agarose (HiMedia, Product no.: MB002) along with SafeDye stain (SRL Chemicals, Product no.: 53261). Gels were run in 1X Tris-Borate EDTA buffer prepared from Tris (Qualigens, Product no.: Q15965), Boric acid (Fisher Scientific, Product no.: 12005) and ethylenediaminetetraacetic acid (EDTA, Fisher Scientific, Product no.: 12635). Oligonucleotides (DNA primers) were obtained from Sigma Aldrich unless specified otherwise and are mentioned in Supplementary Table 1.

### B. Methods

#### Transcription of RNAs

All the RNA sequences are same as used in previous studies Vaidya et al. (2012); Ameta et al. (2021). RNAs were *in vitro* transcribed as described previously Arsène et al. (2018). For transcriptions, the dsDNA templates were prepared by amplifying the wild-type WXYZ template using specific primers. For PCRs, WXYZ template (at 25 *pg/μL*) was mixed with 0.5 *μM* of respective forward and reverse primers (see Supplementary Table 1) in 1X PCR buffer (Thermo Scientific), 0.2 mM dNTPs, 0.02 *U/μL* of polymerase (Thermo Scientific *Taq* polymerase, Product No.: EP402) and amplified using the following protocol: initial denaturation 94°C/5 min, then 28 cycles of denaturing 92°C/1 *min*, annealing at 57°C/1 *min*, extension at 72°C/1 *min* and a final extension of 72°C/5 *min*. PCR products were isopropanol precipitated by adding 1/10^*th*^ volume of 3 *M* sodium acetate pH 5.5 and 1.2 volume of 100% isopropanol to the PCR reaction and centrifuging at 13.8 rcf for 60 *min* at 10°C. Pellets were vacuum dried, re-suspended in 20-30 *μL* of water and used for *in vitro* transcription reactions as described in Arsène et al. (2018). Briefly, the re-suspended PCR products were mixed with 4 mM of each NTPs, 1X Transcription buffer (Thermo Scientific), 12 mM MgCl_2_, 3 *U/μL* of T7 polymerase (lab-made), and incubated at 37°C for 8 *h*. The transcribed RNAs were purified on 12% denaturing polyacrylamide gels, specific bands were eluted from gels, and isopropanol precipitated in the same way as mentioned above. RNA concentrations were measured using NanoDrop 2000 (Thermo Scientific). The Z fragment RNA used here was custom synthesized by IDT (Integrated DNA Technologies, Belgium) and used without further purification.

#### Formation of PAA-spermine coacervate compartments

In general coacervate compartments were prepared in 25 *μL* scale volume (unless specified otherwise) by mixing thoroughly at first 5 mM spermine and 25 mM Tris-HCl (pH 8.0) together in water and then adding 12.5 mM of PAA (HMW). Then, 1X ribozyme reaction buffer (AZ Buffer: 30 mM EPPS, 50 mM MgCl_2_, pH 7.0) was added and solution was mixed again using pipetting. For the RNA reactions, substrate RNA fragments (after folding) were added prior to spermine addition (see Methods below). For the coacervate droplets, these compartments were used directly. For the macrophase, the coacervate were settled by centrifuging the tubes at 100 rcf for 20 *min* at 4°C (Supplementary Fig. S2). After pipetting out the bulk aqueous phase, the remaining coacervates at the bottom of the tube were used as coacervate macrophase. Both, droplets and macrophase were analyzed by microscopy as well FRAP (see below). To facilitate this, additionally a 30nt 5’-Alexa-488 labeled DNA oligonucleotide (custom synthesized from IDT, Integrated DNA Technologies, Belgium, Supplementary Table 1) was added to the coacervate samples, at a 10 *nM* concentration, prior to the addition of Tris and spermine.

#### Fluorescence recovery after photobleaching (FRAP) measurements

The FRAP measurements were carried out both on the coacervate droplets as well as separated coacervate macrophase (Fig. 1C, D), and samples were prepared as described above. For FRAP, a PDMS (polydimethylsiloxane, SYLGARD™ 184, DOW) sample chamber with a glass cover slip was prepared on which ~100 *μL* of sample was added and covered with another glass cover slip. Coacervate droplet samples were kept for 30 *min* to settle down whereas separated macrophase samples were used directly after sealing the glass slides with nail polish from the outside (sandwiching between two glass cover slips). Measurements were carried out on the Olympus FV3000 confocal microscope using cellSens software. For each sample, a region of interest (ROI) was photobleached for 10 *s* using 100% laser power at 488 nm. This is followed by measuring the fluorescence recovery time of ROI. For coacervate droplets (Fig. 2D), ROI circle of diameter 5 *μm* was chosen and the time series was acquired at 1.6 fps for 60 *s*. For separated macrophase samples, the fluorescence recovery time series was acquired at 0.3 fps for 300 *s* with ROI circle diameter of 18 *μm*. The fluorescence intensities from the image sequences were measured using Fiji software (https://imagej.net/Fiji, Schindelin et al. (2012)) to generate the recovery curve. Recovery data was normalized using the equation mentioned below explained by Jia *et al*. Jia et al. (2014)

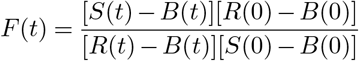

where *F* (*t*) is the normalized fluorescence intensity of the ROI at time *t*, *S*(*t*) is sample ROI intensity, *R*(*t*) is reference ROI intensity, *B*(*t*) is background ROI intensity. The data was fitted using first order exponential equation given below:

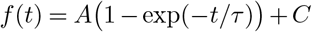

where *f* (*t*) represents the normalized fluorescence intensity, *A* represents the amplitude of the recovery, and *C* represents the y-intercept. By fitting the experimental data in this equation, the half-time *t*_1/2_ = ln(2)*τ* was calculated where, *τ* represents the recovery time.

The apparent diffusion coefficient (*D_app_*) for 2D diffusion was calculated using the equation below Axelrod et al. (1976); Poudyal et al. (2019):

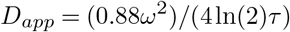

where *ω*, represents the ROI radius.

#### Radioactive labeling of RNAs

To facilitate the measurement of the product formation by gel electrophoresis, all the reaction were doped with minor amounts (0.01 *μM*) of radioactively labeled W or WXY fragment (*γ*^32^*P*). These radioactively labeled RNAs were prepared by mixing 1 *μM* of RNA with 1X of Shrimp Alkaline Phosphatase buffer (rSAP reaction buffer, New England BioLabs), 1 *U/μL* of rSAP enzyme (New England BioLabs, Product no.: M0371S), and water in 10 *μL* reaction scale. Samples were incubated at 37°C for 40 *min* and enzyme was heat inactivated at 70°C for 10 *min*. The SAP reaction samples were then directly used for kinase reaction by mixing 0.5X of polynucleotide kinase buffer (PNK Buffer A, Thermo Scientific), 5 *μL* of *γ*^32^*P* - ATP (10 *mCi*/*mL*, BRIT Hyderabad, India, Product No.: PLC 101), 0.4 *U/μL* PNK enzyme (Thermo Scientific, Product No.: EK0031) and water in 20 *μL* reaction scale. The samples were incubated at 37°C for 1 *h* and reaction was stopped by adding equal volume of gel-loading buffer (90% formamide, 0.01% xylene cyanol, 0.01% bromophenol blue). Labeled RNAs were purified on 12% denaturing polyacrylamide gels, eluted in 0.3 M of sodium acetate pH 5.5 (2 h at 37°C) followed by isopropanol precipitation as mentioned above. The pelleted RNAs were resuspened in water and used for the RNA catalysis experiments.

#### Four fragment self-assembly reactions inside the coacervate compartments

For four fragment assembly in coacervates, 25 *μL* coacervate samples were prepared by mixing substrate RNA fragments (W, X, Y, Z at 0.75 *μM* each) in water, heated at 80°C/3 *min* and then incubated 4°C/3 *min* (on ice) to fold the RNAs. Then, Tris, spermine and PAA were added in the same way and at the same concentrations as described above followed by addition of 1X ribozyme reaction buffer (30 mM EPPS, 50 mM MgCl_2_, pH 7.0). Solution was mixed thoroughly by pipetting. For the macrophase, bulk of the aqueous phase was removed from top and the settled condensed phase at the bottom of the tube was incubated at 48°C (Supplementary Fig. S2). In the case of coacervate droplets, the solution was incubated at first and then to analyze the product formation from the droplets, coacervates were separated from the bulk solution in similar way as mentioned above. In each of the case, for the time-courses, a separate reaction tube was prepared for each time-point. For preparing samples for the gel analysis, after centrifugation and separating the bulk of the solution, water was added to the separated coacervates 25 *μL*, followed by 25 *μL* of PAA-stopping solution (50 mM PAA-LMW, 166 mM EDTA, 333 mM NaOH) and the samples were then vortexed vigorously. Then from this mixture, 20 *μL* was mixed with 20 *μL* of stopping solution (90% formamide, 50 mM EDTA, 0.01% xylene cyanol, 0.01% bromophenol blue) and analyzed on 12% denaturing polyacrylamide gels. For the bulk aqueous phase control, the reactions were carried without any of the coacervate reagents (PAA, Spermine, Tris), however, the time-points were collected and processed in the same way as the coacervate samples.

#### Lipid-vesicle coating of coacervate droplets and catalysis

The small unilamellar vesicles (SUVs) were prepared by using a previously reported protocol Kumar et al. (2022). Briefly, 25 *μL* of DOPC (1,2-dioleoyl-sn-glycero-3-phosphocholine, 25 *mg*/*mL* prepared in chloroform; Avanti Polar Lipids) and 5 *μL* of Liss-Rhod-PE (1,2-dioleoyl-sn-glycero-3-phosphoethanolamine-N-(lissamine rhodamine B sulfonyl), 10 *μg/mL* prepared in chloroform; Avanti Polar Lipids) were mixed in a 2 *mL* eppendorf tube, vacuum-dried overnight followed by drying under Nitrogen before resuspended in 1 *mL* nuclease-free water. To prepare SUVs, a probe sonicator (VC 750 (750W), stepped microtip with diameter 3 mm) with an on/off pulse of 1.5 *s*/1.5 *s* and 32% of the maximum sonicator amplitude were used. The tube was placed in an ice bath in between the sonication steps (sonication for 30 *s* with a 1 *min* gap in between). Then the tube was briefly spinned (using micro-centrifuge) and SUVs were stored at 4°C. To carry out four-fragment assembly of WXYZ catalyst inside the vesicle-coated coacervate droplets, at first, a coacervate solution was prepared as mentioned above by adding all the four RNA fragments (W, X, Y, Z) at 0.75 *μM* each (after folding together), 5 mM spermine and 25 mM Tris-HCl (pH 8.0) in water. Then 12.5 mM of PAA (HMW) was added, solution was mixed thoroughly by pipetting followed by addition 1X ribozyme reaction buffer (AZ Buffer: 30 mM EPPS, 50 mM MgCl_2_, pH 7.0) and again mixing by pipetting. Finally, 4 *μL* of SUVs solution was added to 25 *μL* coacervate solution, mixed by pipetting, and kept on ice for 5 *min* to allow for coating. Then to initiate reaction, the solution was incubated at 48°C. As mentioned above, a separate reaction tube was prepared for each time-point and samples were processed in the same way without any additional steps.

To measure the stability of vesicle-coated coacervate droplets, their size distribution was analyzed over the time. Lipid-vesicle coated coacervate droplets were prepared in the same way as mentioned above, incubated at 48°C, and analyzed under the microscope. The images were captured in the fluorescent channel of inverted microscope (100x oil objective, exposure time 100 *msec*) and the droplet sizes were calculated using the *analyse particle* function in Fiji software (https://imagej.net/software/fiji/). To analyze the leakiness or diffusion of oligonucleotide from the bulk phase into vesicle-coated coacervate droplets, droplets were prepared with a 30nt long fluorescently labelled DNA oligonucleotide (Alexa-488 labeled, same as used in Fig. 1, see Supplementary Table 1), both alone (Fig. 2D), *top*) as well as hybridized to the ~200nt long WXYZ RNA (Fig. 2D), *bottom*), and then coated with DOPC vesicles (without doping with Liss-Rhod-PE in order to avoid bleed through of fluorescence signal). Then sample was incubated at 48°C and images were captured in the fluorescent channel of inverted microscope (100x oil objective, exposure time 100 *ms*) at 0 *h* and at 4 *h*. The fluorescent images were analysed using Fiji software (https://imagej.net/software/fiji/) for the change in fluorescence intensity inside the vesicle-coated coacervate droplets. To analyze the images, a line was taken through the centre of the droplet on the raw image (without any pre-processing of the images) and fluorescence intensity profile was extracted along that line. For each sample and the time-point, ~10 droplets of approximately similar sizes were analyzed and the average fluorescence intensity of several droplets as a function of distance from the centre of the droplet was plotted (Fig. 2D).

#### Kinetic Modelling for the WXYZ assembly

The sigmoidal nature of the time course of the self-assembly of *Azoarcus* ribozyme (WXYZ) from the four fragments (W, X, Y, and Z) was confirmed by analyzing the time-course data with a simple kinetic model. In the chemical reaction model, which is reduced from the full detailed model described in Hayden *et al*. Hayden and Lehman (2006); Hayden et al. (2008), we assumed two types of reactions:

1. A instantaneous non-covalent complex (W:X:Y:Z) formation from the four fragments (W, X, Y, and Z),

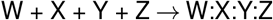

whose association rate (estimated as ~ 10^2^ − 10^3^ min^−1^ for 1*μ*M of fragments, in the previous studies under similar condition Vaidya et al. (2012); Hayden et al. (2008); Arsène et al. (2018)) is much faster than the time scale of the measurements. While its dissociation rate is much slower (estimated as ~ 10^−2^ min^−1^ Vaidya et al. (2012)), thus the fraction of the fragments forming the complex is saturated, since the dissociation constant is far smaller than the concentration of the fragments ~ 1*μ*M.
2. Reversible covalent formation,

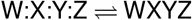

is catalyzed both by W:X:Y:Z and WXYZ with rates *k_a_* and *k_b_*, respectively. The ratio between the forward and the backward reactions is 1 : *δ*.

Here, we also assume the four fragments (W, X, Y, and Z) and the ribozyme (WXYZ) do not decay during the observation. In the experiment, we set the equal molar of fragments as the initial condition. Then, defining the initial fraction of each fragment as unity, the sum of the fraction of W:X:Y:Z and WXYZ is conserved as unity since the complex formation is immediate, and all of the fragments exist as either W:X:Y:Z or WXYZ.

Under the above assumptions, the model rate equation is given:

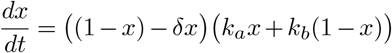

where 1 − *x* and *x* are the fraction of W:X:Y:Z and WXYZ, respectively. This can be simply transformed into the logistic differential equation,

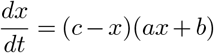

where *c* = 1/(1 + *δ*), *a* = (1 + *δ*)(*k_a_* − *k_b_*), and *b* = (1 + *δ*)*k_b_*. The solution for *x* is given as

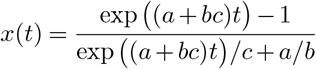

We fitted t he f unction w ith t he t ime c ourse d ata obtained by the gel electrophoresis data (Fig. 2) by minimizing the sum of weighted squared residuals Press et al. (2007). Note that, as a result of the fitting, *a* > 0 (i.e., *k_a_* > *k_b_*) indicates the evidence of autocatalytic reaction (Supplementary Table 2), i.e., the product ribozyme formation as a result of positive feedback from the product itself.

#### Measuring autocatalytic rate-constants inside the coacervate macrophase

Effect of catalyst and autocatalytic rate constants inside the coacervates were measured using the same strategy as developed by von Kiedrowski von Kiedrowski (1993), and used for the bulk reactions in the previous studies Yeates et al. (2016); Ameta et al. (2021) by analyzing two-fragment self-assembly reaction (between WXY and Z RNAs). However, here low concentration of substrate RNAs and WXYZ catalyst were used. Briefly, coacervate samples were prepared as described above using 0.1 *μM* of WXY (along with 0.01 *μM* ^32^*P* radioactive labeled WXY) and Z RNAs each along with different concentration of seeded WXYZ catalyst (at 0, 0.015, 0.15, 0.3, 0.5 *μM*, having same IGS as WXY fragment). To measure initial rate of reaction, samples were incubated for very short period of time (~ 12 *min*) and reactions were stopped to prepare gel samples in the same way as described above. Initial rates of formation of covalent WXYZ were derived from the time courses of assembly of WXY and Z fragments (only from the linear portion of the time-courses) in presence of different concentration of seeded product (WXYZ) catalyst added at the beginning of the reaction. As observed earlier Yeates et al. (2016); Ameta et al. (2021), the initial rate of formation of WXYZ (*r*_0_) was found to depend linearly on the concentration of the seeded WXYZ. This dependence can be described using the following relationship:

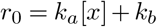

where *x* is the concentration of the covalent ribozyme, *k_a_* is obtained from the slope and *k_b_* is obtained from the y-intercept. Here, *k_a_* quantifies the formation of WXYZ product synthesis by covalent WXYZ ribozyme and *k_b_* quantifies the formation of WXYZ product synthesis by non-covalent ribozyme assembly (trans-catalysis by non-covalent complex of WXY and Z fragments Hayden et al. (2008); Vaidya et al. (2012)). For the aqueous phase control (bulk reaction without any polymers), due to low reaction rates in absence of coacervation, self-assembly reactions were carried for longer time period (up to 90 *min*) and higher concentration of WXYZ catalyst (at 0, 0.5, 0.75, 1.0, 2.0 *μM* were added at the beginning of each reaction. Rest of the processes were done in a similar manner as for the coacervate samples.

#### Network formation inside the coacervate compartments

To confirm the formation of network inside the coacervates, respective RNA fragments to construct two-, three- and four nodes networks were mixed together at 0.5 *μM* each along with equivalent amount of Z RNA fragment, and 0.01 *μM* of ^32^*P*-labeled WXY RNA fragment for UU in water at 25 *μL* scale. The samples were then heated at 80°C/3 min and then incubated 4°C/3 *min* (on ice) to fold the RNAs and coacervation was induced as described above. The coacervate droplets were directly incubated at 48°C (for 1 *h*) and then separated post-incubation for the polyacrylamide gel analysis whereas the coacervate macrophase was separated from bulk of the solution (as above, Supplementary Fig. S2) and incubated at 48°C for 1 *h*. Timepoints and gel analysis was done in the same way as described above in four-fragment assembly section.

#### Network composition measurements

To measure the network composition inside coacervates, WXY RNA fragments for the respective nodes at 0.5 *μM* each along with equivalent amount of Z RNA fragment were used. For measuring amount of different nodes from the same network, multiple reactions (equal to the number of nodes to be measured) were set-up in parallel where each one is doped with 0.01 *μM* of the respective ^32^*P* - labeled WXY RNA fragment. For example, in the case of two nodes network (UU, AA), two parallel reactions were set-up where one reaction was doped with ^32^*P* - labeled UU WXY RNA fragment and another one was doped with ^32^*P*-labeled AA WXY RNA fragment. After adding all the RNAs, the samples were then heated at 80°C/3 *min* and then incubated 4°C/3 *min* (on ice) to fold the RNAs, followed by inducing coacervation as described above. The coacervate droplets were directly incubated at 48°C (for 30 *min*) and then separated postincubation for the polyacrylamide gel analysis whereas the macrophase was separated from bulk of the solution (as above) and incubated at 48°C for 30 *min*. Time-points and gel analysis was done in the same way as described above in four-fragment assembly section.

#### Network perturbation

In order to study network perturbation, at first coacervate solution containing WXY fragments of UA and UG nodes at 0.5 *μM* each along with equivalent amount of Z RNA fragment was prepared in a similar way as mentioned in the section above (Network composition). For the droplet phase perturbation, a separate coacervate droplet population containing 0.5 *μ*M of WXY fragment for CA along with 0.5 *μ*M of Z RNA fragments was prepared and mixed with coacervate droplets containing UA and UG. The solution was mixed well by pipetting and incubated at 48°C. For the perturbation in coacervate macrophase, after separating the condensed phase containing the WXY fragments of UA and UG from bulk of the solution, 0.5 *μM* of WXY fragment for CA along with 0.5 *μM* of Z RNA fragments were added on the top of the coacervate macrophase (~ 4 *μL*)and then samples were incubated 48°C. For aqueous phase control, the reactions were carried out without any of the coacervate reagents (PAA, spermine, Tris-buffer) however, the time-points were collected and processed in the same way as the coacervate samples.

## Acknowledgments

We thank Philippe Nghe, Sandeep Krishna and Madan Rao for discussions and comments on the manuscript. We thank Pranav Sharma and Anupam Singh for help with experiments, the laboratories of Arati Ramesh for providing T7 RNA polymerase, and of Praveen Vemula for providing high molecular weight poly(acrylic acid). We acknowledge access to the radiation, instrumentation and central imaging facilities at the NCBS, Bangalore.

## Author Contributions

S.T. and S.A. conceived the idea. S.T., S.A. and M.K. designed the study. S.A., M.K., N.C., P.S. and D.G, performed the experiments and analyzed the data. Y.J.M. performed the kinetic modelling. S.T. and S.A. wrote the paper.

## Data Availability

The data that support the findings of this study are available from the corresponding authors upon reasonable request.

## Code Availability

Not applicable

## Supplementary Information

### Text 1: Autocatalytic nature of the WXYZ assembly and effect of phase-separation

*Azoarcus* ribozyme self-assembles from its fragments (two, three or four) in an autocatalytic manner Hayden and Lehman (2006); Hayden et al. (2008). The autocatalytic nature of the reaction can be observed by plotting the amount of covalently self-assembled product formed as a function of time (as observed in Fig. 2). Furthermore, we have also observed the positive effect of catalyst on the four-fragment assembly of the *Azoarcus* ribozyme (Supplementary Fig. S4). Though the sigmoidal curve can be easily observed for the four-fragment assembly Hayden and Lehman (2006), it is not so straightforward for the two-fragment assembly, owing to its faster reaction timescale Hayden et al. (2008); Vaidya et al. (2012); Ameta et al. (2021). This can somewhat be resolved by slowing down the reaction either by lowering the concentration of substrate or magnesium. Even though the sigmoidal nature of the curve is a consequence of autocatalytic nature of the reaction, a better demonstration is through the seeding experiments, where substrate of fixed concentration is seeded with varying catalyst concentrations (product of the reaction) added at the starting of the reaction von Kiedrowski (1993). In such an experiment, the positive feedback is confirmed if there is an increase in the rate of reaction as a function of (seed) catalyst concentration. We have observed a clear positive effect of the catalyst in the seeding experiment for the assembly of the *Azoarcus* ribozyme (Supplementary Fig. S6). In addition, such measurements also allow to quantify autocatalytic rate constants, *k_a_* (feedback from the product of the reaction, *i.e*. covalent WXYZ catalyst) and *k_b_* (background reaction, *i.e*. catalysis by non-covalent WXYZ complexes). Please note that the catalysis by non-covalent complexes is due to the formation of active catalytic pocket formed even in the fragmented version of the ribozyme Hayden et al. (2008); Ameta et al. (2021) and not due to the spontaneous background (uncatalyzed). While in literature von Kiedrowski (1993); Ameta et al. (2021a), *k_b_* accounts for the reaction carried out due to the spontaneous reaction, for simplicity, in the case of *Azoarcus* ribozyme assembly we use *k_b_* for the catalysis by non-covalent WXYZ complexes. Therefore in the case of *Azoarcus* ribozyme, there are two catalytic components – covalent WXYZ and non-covalent complexes. In the initial phase of the reaction, due to the higher concentration of RNA fragments, the non-covalent complexes form really fast (<1 *min*) and majorly govern the self-assembly process Vaidya et al. (2012); Ameta et al. (2021). This is more evident in the phase-separated coacervate compartments as they tend to concentrate biomolecules inside them (Supplementary Fig. S8) such that the effect of *k_b_* is more pronounced on the assembly process of the *Azoarcus* ribozyme. This is also captured in our measurements (Supplementary Table 2 and Fig. S6), and as a consequence, the formation of autocatalytic reaction networks is indeed affected by the catalytic activity of non-covalent complexes, *k_b_* in the phase-separated compartment rather than *k_a_*.

### Text 2: Perturbation scenario in coacervate compartments

Studying the perturbation scenarios in self-replicating chemistries is crucial in developing an evolvable protocell as one of the significant roles of compartment is to protect the identity (of the replicating system). The network composition are indeed amenable to the perturbations which can occur due to spatial and temporal heterogeneity of the prebiotic environment, or due to the presence of parasitic/competing species. Furthermore, it has been (theoretically) shown that slight exchange between compartments or with the environment may even aid in the maintenance of self-replicating systems, particularly ribozyme formation using substrate fragments (Kamimura A. *et al*. Kamimura et al. (2019)). While the notion of parasitic species is not clear in the autocatalytic reaction networks, specially the cooperative networks formed by *Azoarcus* ribozymes Vaidya et al. (2012), perturbation due to environmental factor or presence of competing catalyst can be conveniently studied.

In the current study, we explore one such scenario where the presence of coacervate compartment render robustness to the network composition (therefore *compositional identity*) against a competing catalyst (Fig. 4). The perturbation experiment is designed to mimic a situation in which a network (with node UA and UG) is already inside the coacervate compartment and is confronted with a competing species (CA) from the environment which targets the isolated node (UG). Without perturbation, UA dominates over UG as there is no node catalyzing the synthesis of UG (apart from weak G→U wobble link Yeates et al. (2016). While in the bulk condition, the presence of CA perturbation changes the ranking of UA and UG, the network composition is found to be preserved in both coacervate droplets as well as macrophase (Fig. 4). Indeed due to the membraneless nature of coacervates, the CA would ultimately diffuse into the compartments and start perturbing the network composition (Supplementary Fig. S11 at sufficiently longer timescales. Such experiments highlight that there is a timescale regime under which even membraneless phase-separated coacervate compartments can provide robustness to the networks.

**Table 1.**
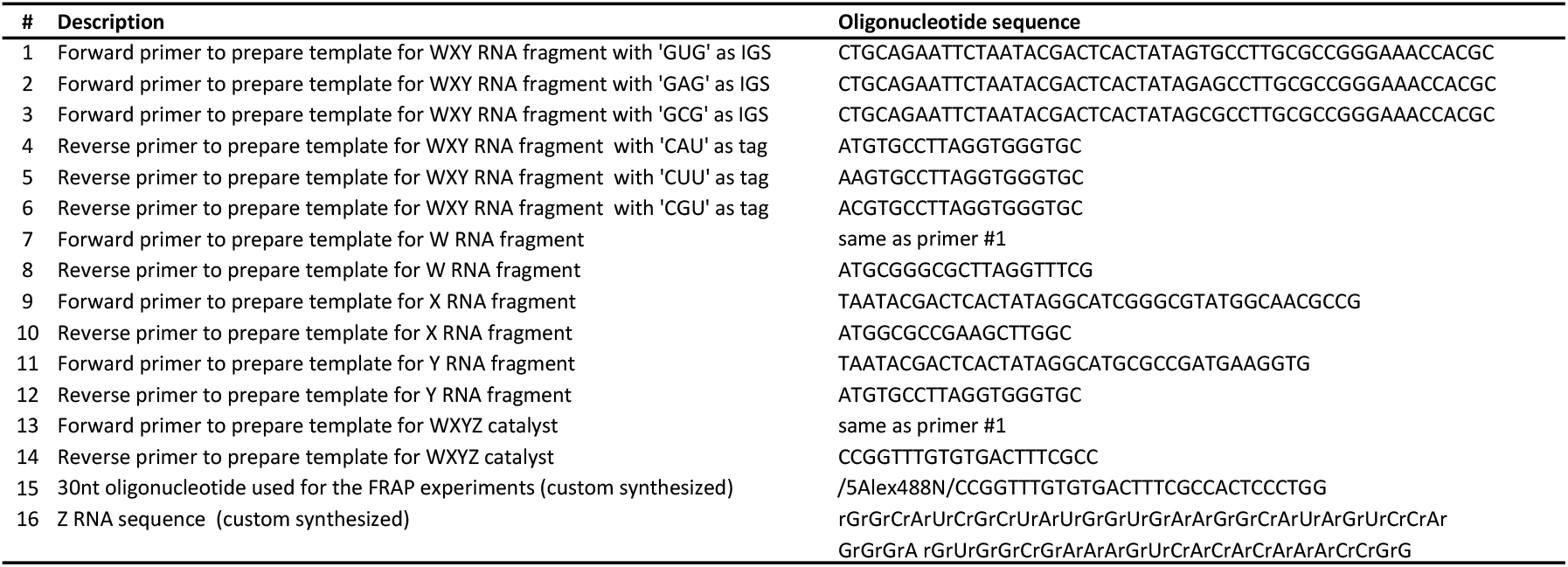
Sequences of all the oligonucleotides used in the study.

**Figure S1.**
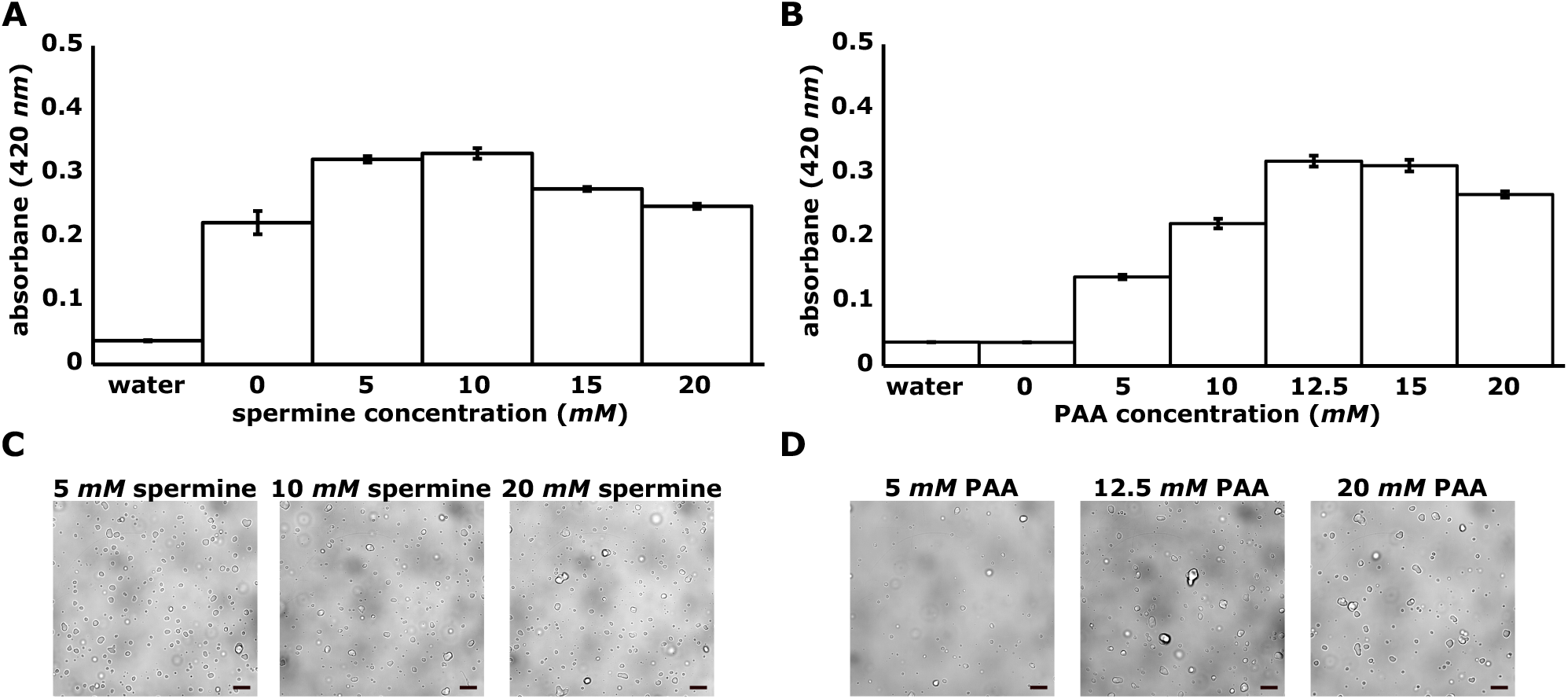
Turbidity assay. Bar graph showing turbidity measured for coacervate solutions at different polymer concentrations to know the dynamic range of coacervation process. Turbidity was quantified by measuring the absorbance of the solution at 420 *nm* wavelength. **A**. At fix concentration of PAA (12.5 mM) and varying spermine concentrations. **B**. At fix concentration of spermine (5 mM) and varying PAA concentrations. All the other conditions of coacervation were kept same as mentioned in Material and Methods. **C, D**. Bright-field microscopy images of PAA-spermine coacervate droplets’ samples from the turbidity assay. C. At fix concentration of PAA (12.5 mM) and varying spermine concentrations (5, 10, 20 mM). D. At fix concentration of spermine (5 mM) and varying PAA concentrations (5, 12.5, 20 mM). Scale bar is 10 *μm* for all the images.

**Figure S2.**
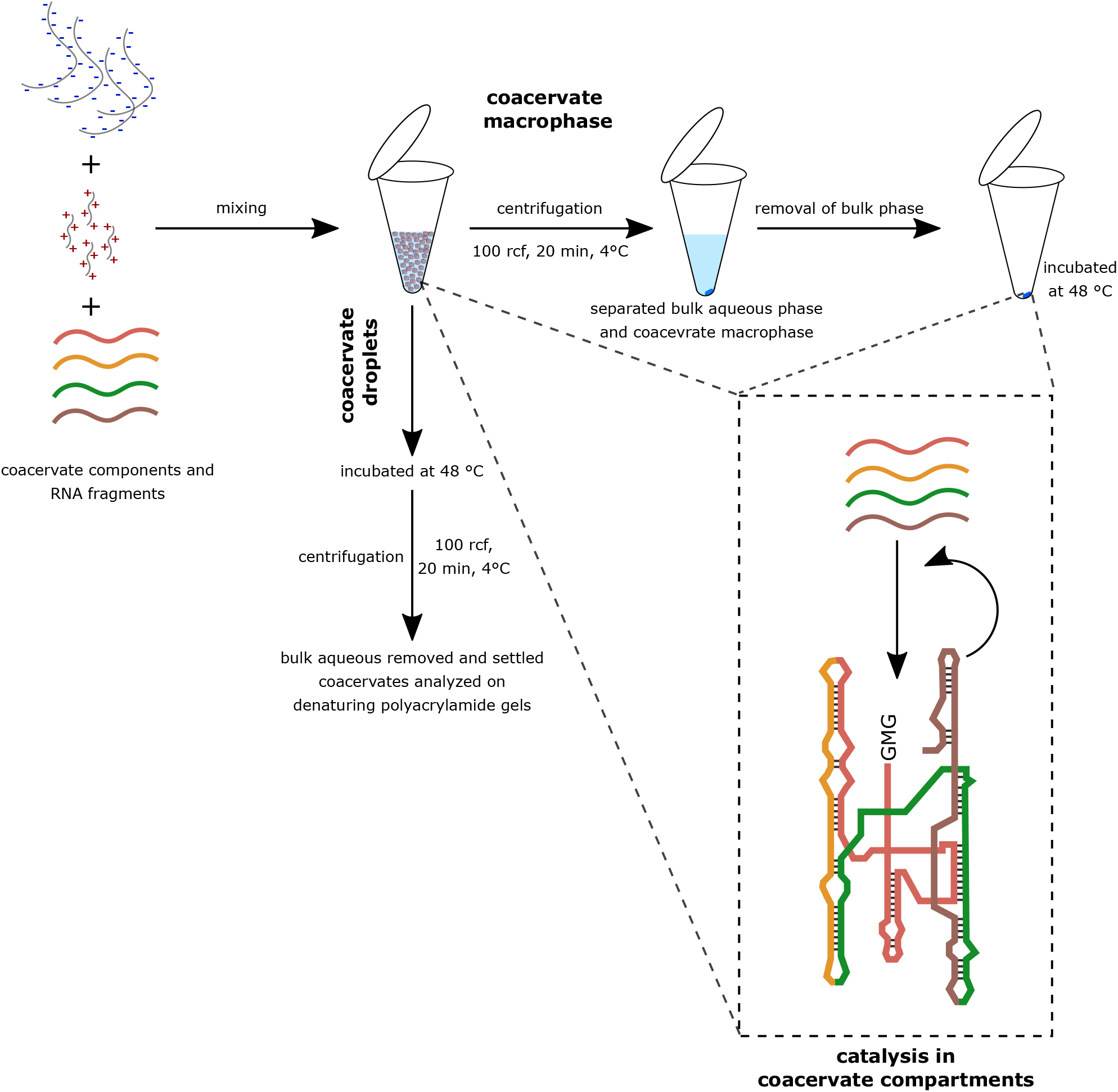
Protocol for analyzing the WXYZ assembly in the coacervate compartments. Schematic of the protocol showing the steps to prepare coacervates and measuring catalysis inside the coacervate droplets and coacervate macrophase. Briefly, all the components of forming coacervates including RNAs are mixed together thoroughly (see Material and Methods) to form coacervates. For the catalysis inside coacervate droplets, the solution was directly incubated at 48°C and then after the reaction, coacervate phase was separated from bulk of the solution by a brief centrifugation (100 rcf, 20 min), the bulk phase was discarded, and the settled coacervate phase was processed for the gel analysis. For the catalysis in coacervate macrophase, the centrifugation was done prior to the incubation and after removing the bulk liquid phase, the settled coacervate macrophase at the bottom of the tube was incubated at 48°C, and then processed for the gel analysis in the same method as described for droplets.

**Figure S3.**
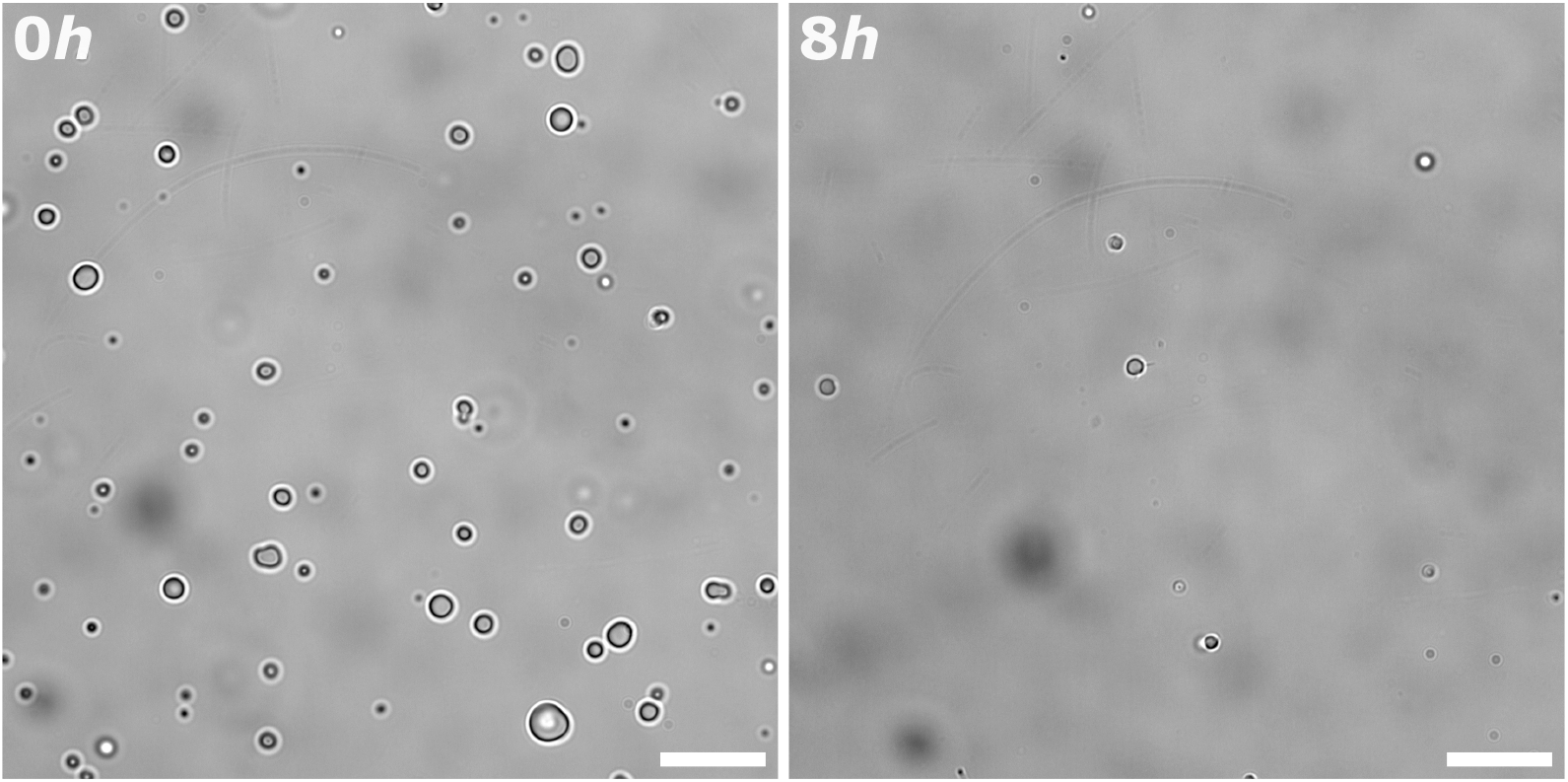
Coacervate droplet stability. Bright-field microscopy images of coacervate droplets (without vesicle coating) to demonstrate that droplets are stable even for longer time period. Image at 0 *h* (*left* and image at 8 *h* (*right*). The result shows that droplets are stable and visible even after incubation at 48°C for longer periods. The density of droplets decreased as they settled down at the bottom of the tube. Scale bar is 15 *μm* for both the images.

**Figure S4.**
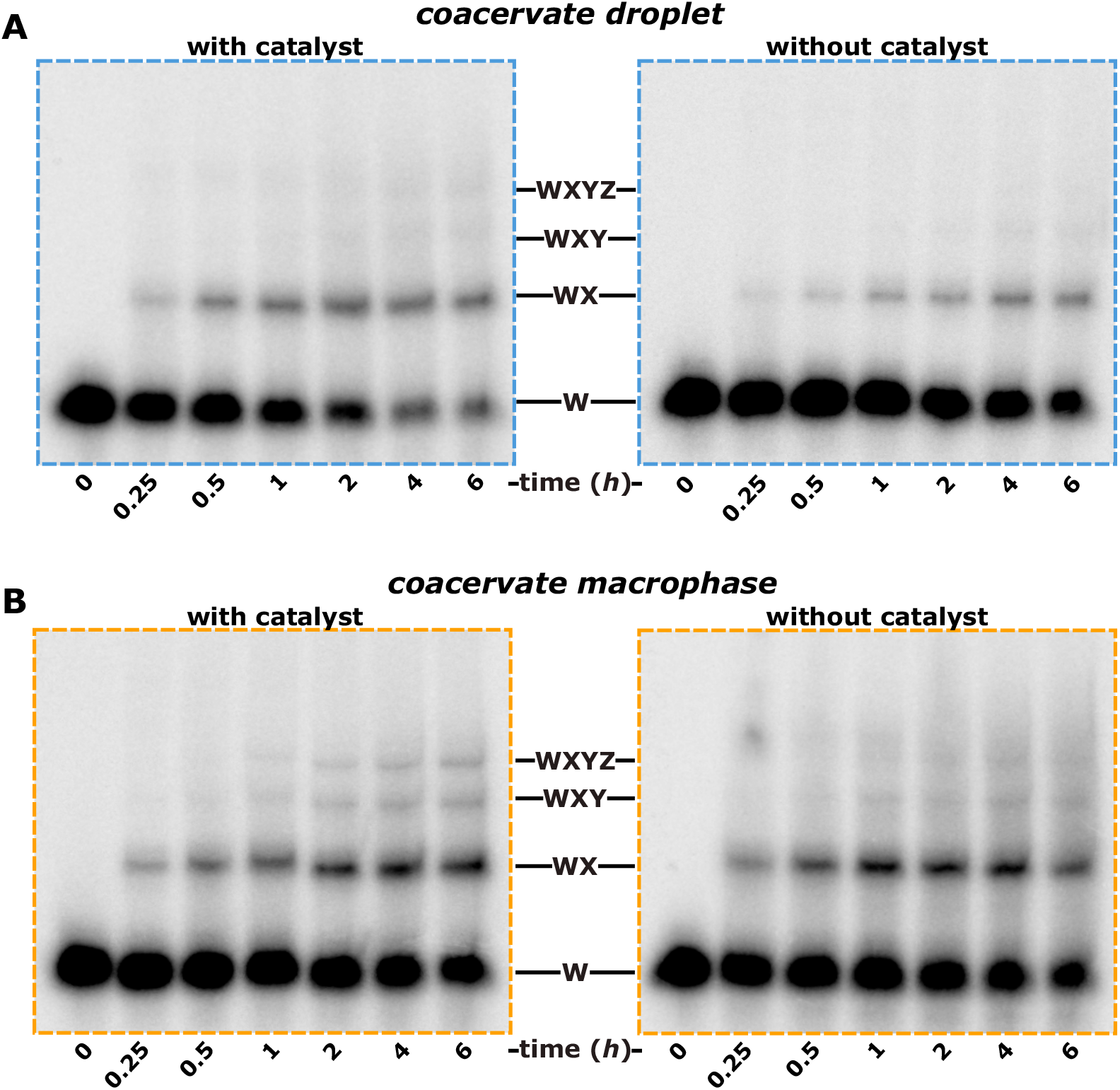
Positive feedback from the product of the reaction on the assembly of WXYZ. Polyacrylamide gel images showing the effect of catalyst (covalent ribozyme, product of the reaction) on its own assembly from the inactive RNA fragments W, X, Y, and Z inside coacervate droplets (**A**) as well as macrophase (**B**). For both the case, assembly is carried out in the presence (*left*) and absence (*right*) of the catalyst (full-length covalent WXYZ ribozyme, 0.5 *μM*). Catalyst is added together with the RNA fragments (0.1 *μM* each) at the start of the reaction. Please see Material and Methods for the experimental detail.

**Table 2.** Kinetic modelling parameters.

| Conditions | a (h<sup>-1</sup>) | b (h<sup>-1</sup>) | c | k<sub>a</sub> (min<sup>-1</sup>) | k<sub>b</sub> (nM min<sup>-1</sup>) |
| --- | --- | --- | --- | --- | --- |
| Bulk reaction | 1.0 ± 0.1 | 0.0020 ± 0.0006 | 0.16 ± 0.01 | 0.0027 ± 0.0003 | 0.00057 ± 0.00017 |
| Coacervate droplets | 0.96 ± 0.25 | 0.010 ± 0.003 | 0.13 ± 0.02 | 0.0021 ± 0.0006 | 0.0022 ± 0.0008 |
| Coacervate macrophase | 1.2 ± 0.3 | 0.03 ± 0.01 | 0.078 ± 0.006 | 0.0015 ± 0.0004 | 0.004 ± 0.001 |
| Coacervate droplets (fitting up to 2h) | 3.4 ± 0.9 | 0.003 ± 0.002 | 0.053 ± 0.007 | 0.0030 ± 0.0009 | 0.0003 ± 0.0002 |
| Coacervate macrophase (fitting up to 2h) | 2.9 ± 0.7 | 0.015 ± 0.005 | 0.056 ± 0.003 | 0.0027 ± 0.0006 | 0.0014 ± 0.0005 |

**Figure S5.**
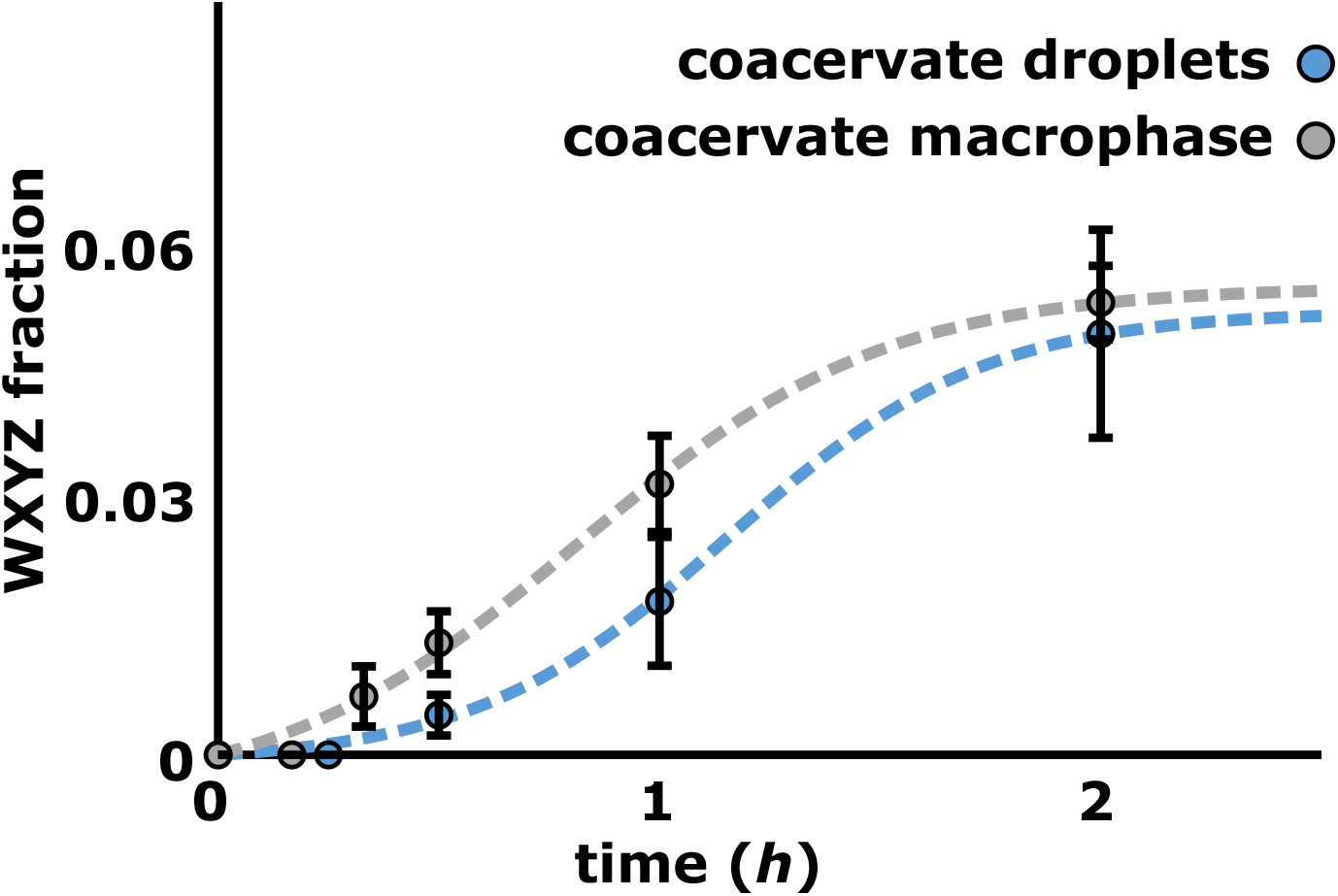
Kinetic modelling fit for the early part of the reaction.

**Figure S6.**
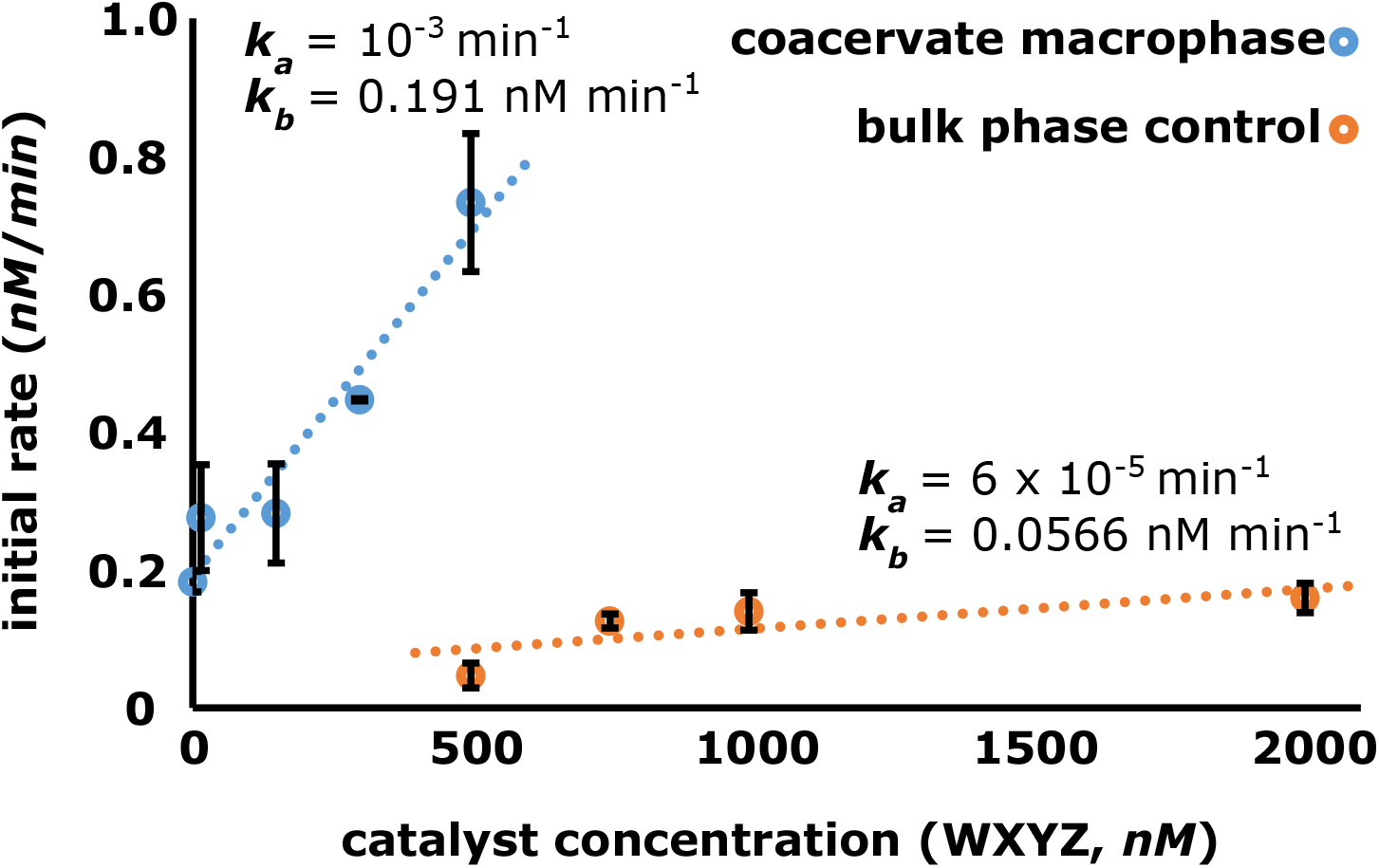
Positive feedback from the product of the reaction. Graph showing the dependence of initial rate of WXYZ formation on the different amount of WXYZ catalyst (product of the reaction) seeded at the start of the reaction. Each data-point on the graph represents the initial rate derived from the time-courses in presence of the different amount of WXYZ. The autocatalytic rate constants are derived in the same way as reported earlier Hayden et al. (2008); Yeates et al. (2016); Ameta et al. (2021). The dashed line in the data points represents a linear fit with *r*^2^ of 0.94 for coacervate macrophase and 0.59 for the bulk phase control. Derived *k*_a_ and *k*_b_ are shown in the inset

**Figure S7.**
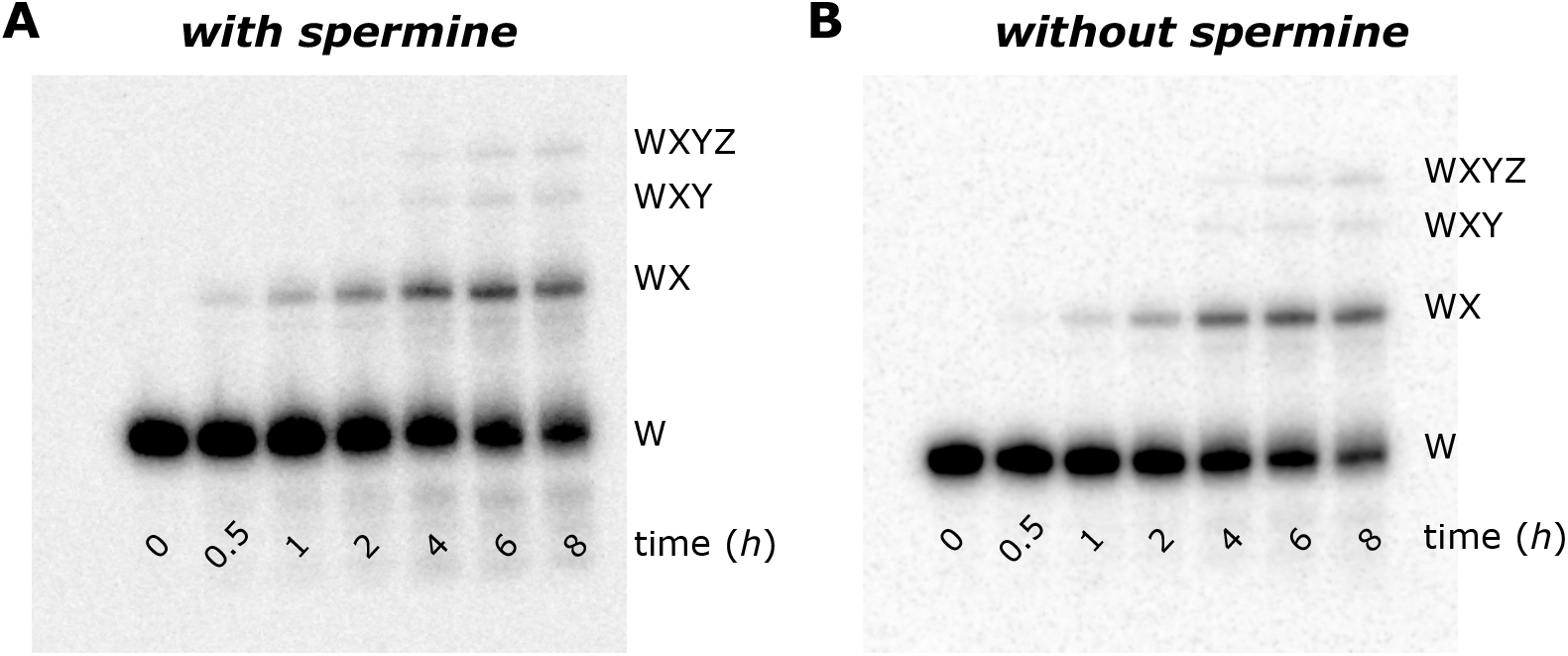
No effect of spermine on self-assembly of WXYZ. Polyacrylamide gel images showing that spermine has no effect on the self-assembly of substrate RNA fragments (W, X, Y, Z) to form WXYZ ribozymes. Here self-assembly reactions are carried out with 0.75 *μM* each W, X, Y and Z RNA fragments in presence of 5 mM spermine (**A**) and in absence of spermine (**B**).

**Figure S8.**
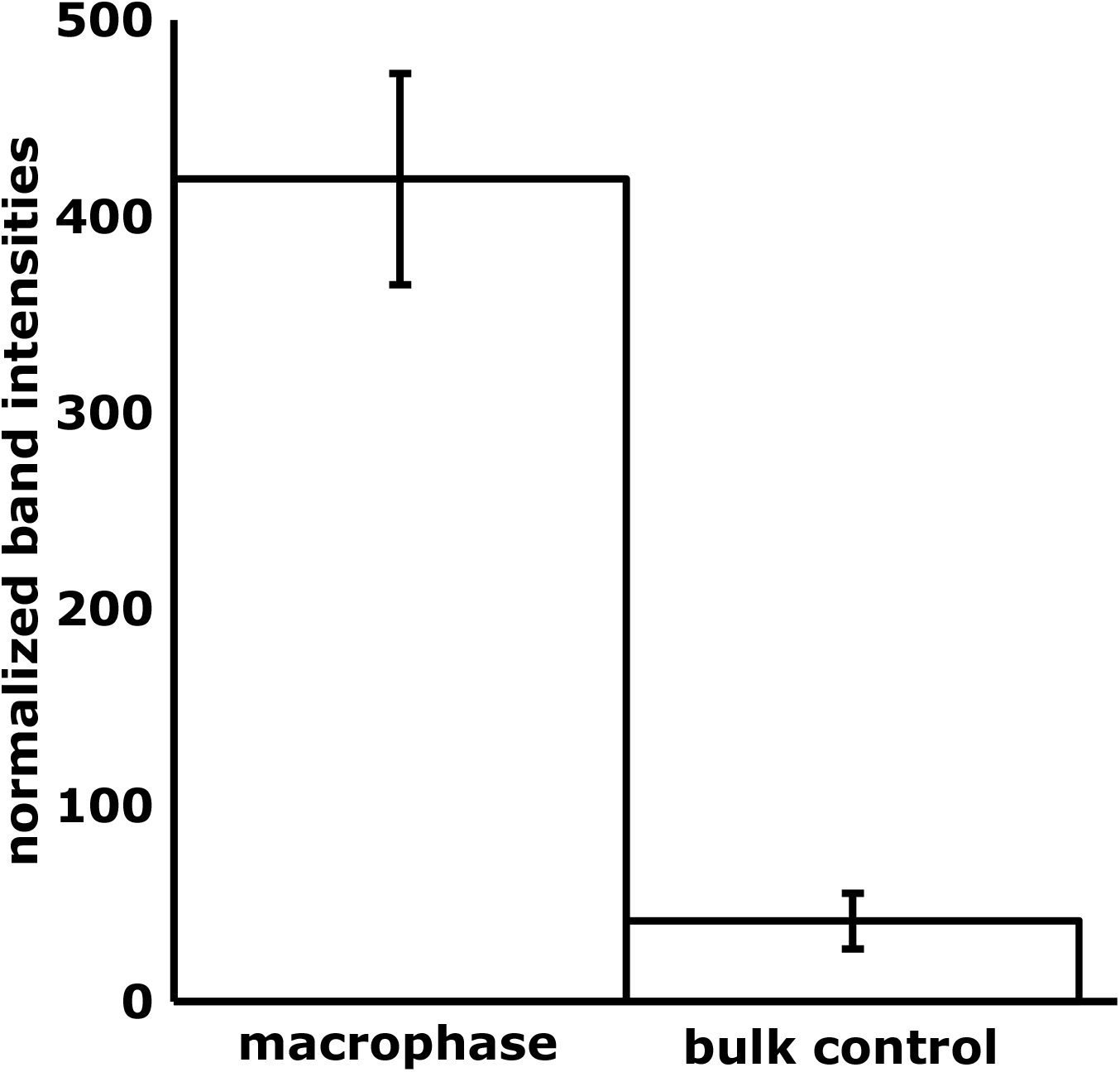
Increase in the RNA amount inside the coacervate macrophase. Bar graph showing the increase in RNA amount inside the coacervate macrophase (~ 50-fold) compared to the bulk phase. To measure the enhancement, samples containing W, X, Y, and Z (0.75 *μM* each) RNA fragments along with spiked amount of radioactive ^32^*P*-labeled W fragment (0.01 *μM*) were prepared with polymers and separated as mentioned above. The bulk as well as coacervate phase were then analyzed on polyacrylamide gels and normalized band intensities are plotted here. For the analysis, settled coacervate macrophase was resuspended in the same amount of water as the bulk sample volume. Then equal volume of both (resuspended coacervate macrophase and bulk control) the samples were loaded on the gel (after adding same amount of PAA stopping and gel loading solutions, see Methods section). After extracting the band intensities, coacervate macrophase band intensities were multiplied by the volume of the water added to equalize the sample volume. All the measurements are done in triplicates and mean is plotted along with standard deviation. The results indicate that there is ~50-fold increase of RNA concentration inside the coacervate corresponding to 37.5 *μM* of RNA (effective) concentration.

**Figure S9.**
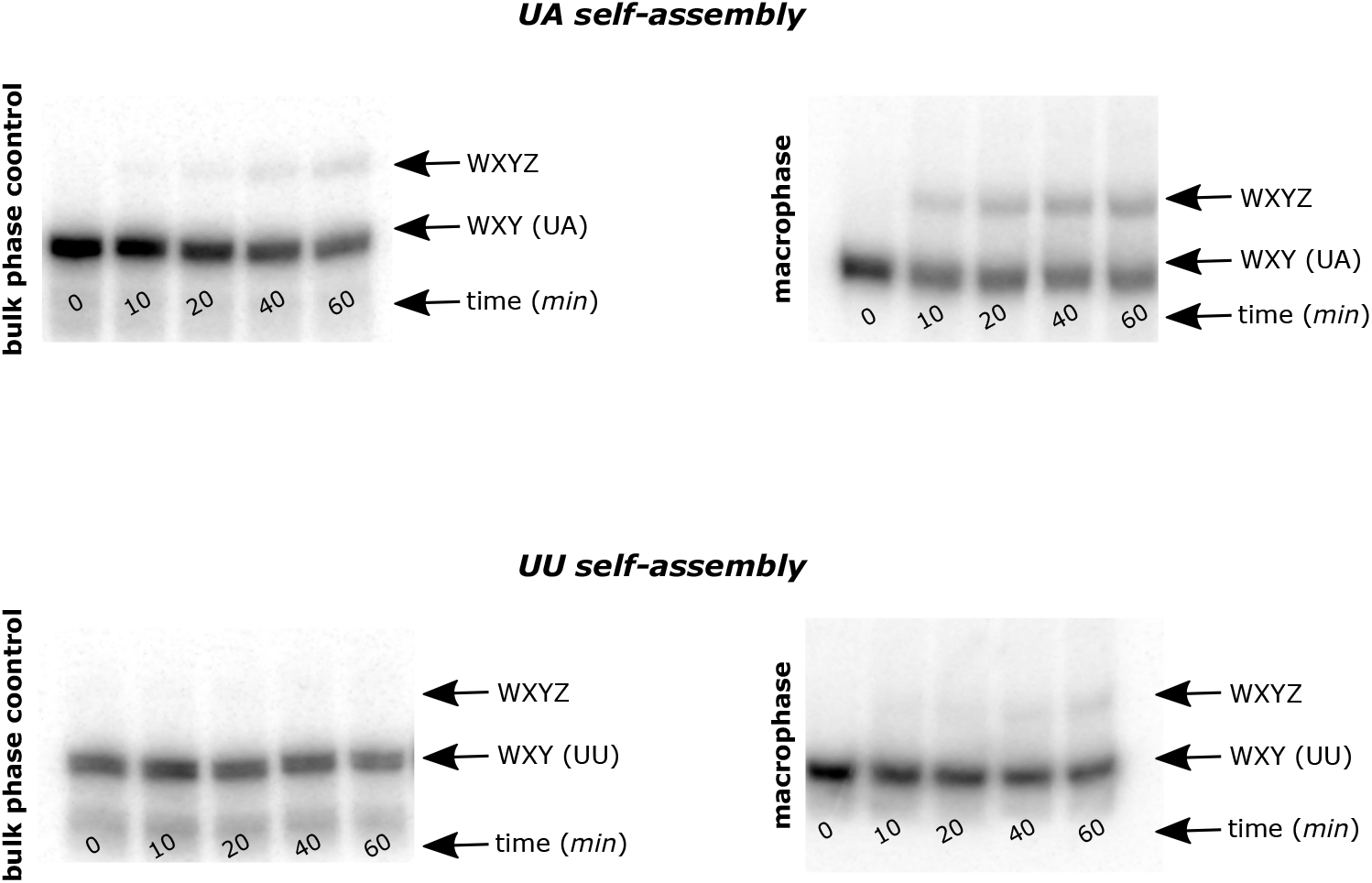
Poor and good self-assembling WXYZ ribozyme inside the coacervate macrophase. Gel pictures showing the product formation for a good self-assembler (UA) in bulk (*top left*) and in the coacervate macrophase (*top right*). The product formation is significantly lower for a poor self-assembler (UU) in bulk (*bottom left*) and even in condensed coacervate macrophase (*bottom right*). Here, two-fragment assembly is carried out between WXY and Z fragments (0.5 *μM* each) of UA and UU in presence of spike amount of respective radioactive ^32^*P*-labeled WXY RNA fragment (0.01 *μM*). Samples are analyzed on a 12% denaturing polyacrylamide gels (see Material and Methods).

**Figure S10.**
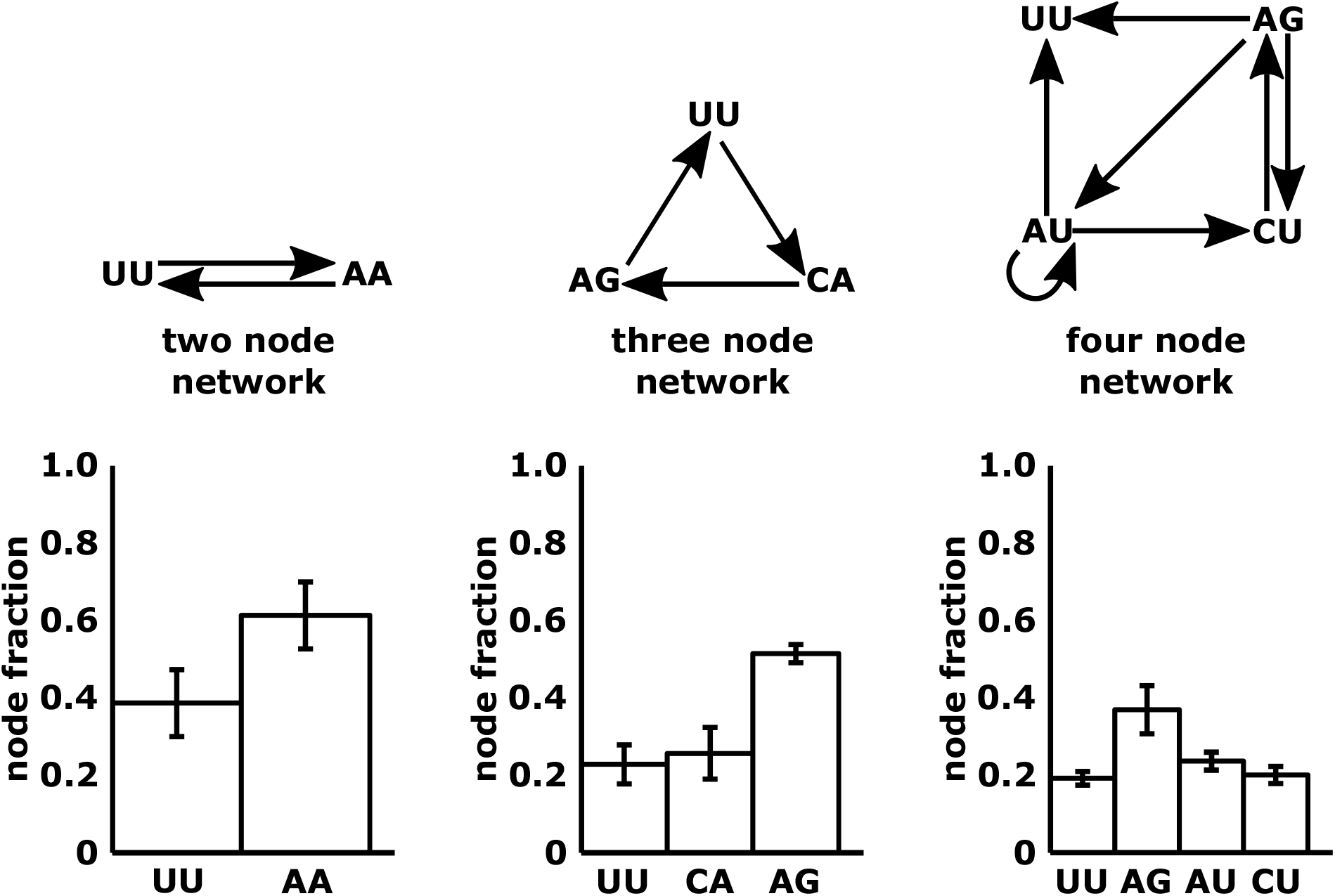
Network compositions of two, three, and four nodes network in absence of any polymer in aqueous phase. Network structures are shown on *top*. Bar-graphs showing the measured composition (*i.e*., relative fraction of individual ribozyme species in the network) of the networks in bulk aqueous phase control for two-node (*left column*), three-node (*middle column*), and four-node networks (*right column*). All the measurements are done in triplicates and mean WXYZ product formation is plotted as fraction w.r.t. to each node along with standard deviation. See Material and Methods for the experimental procedure.

**Figure S11.**
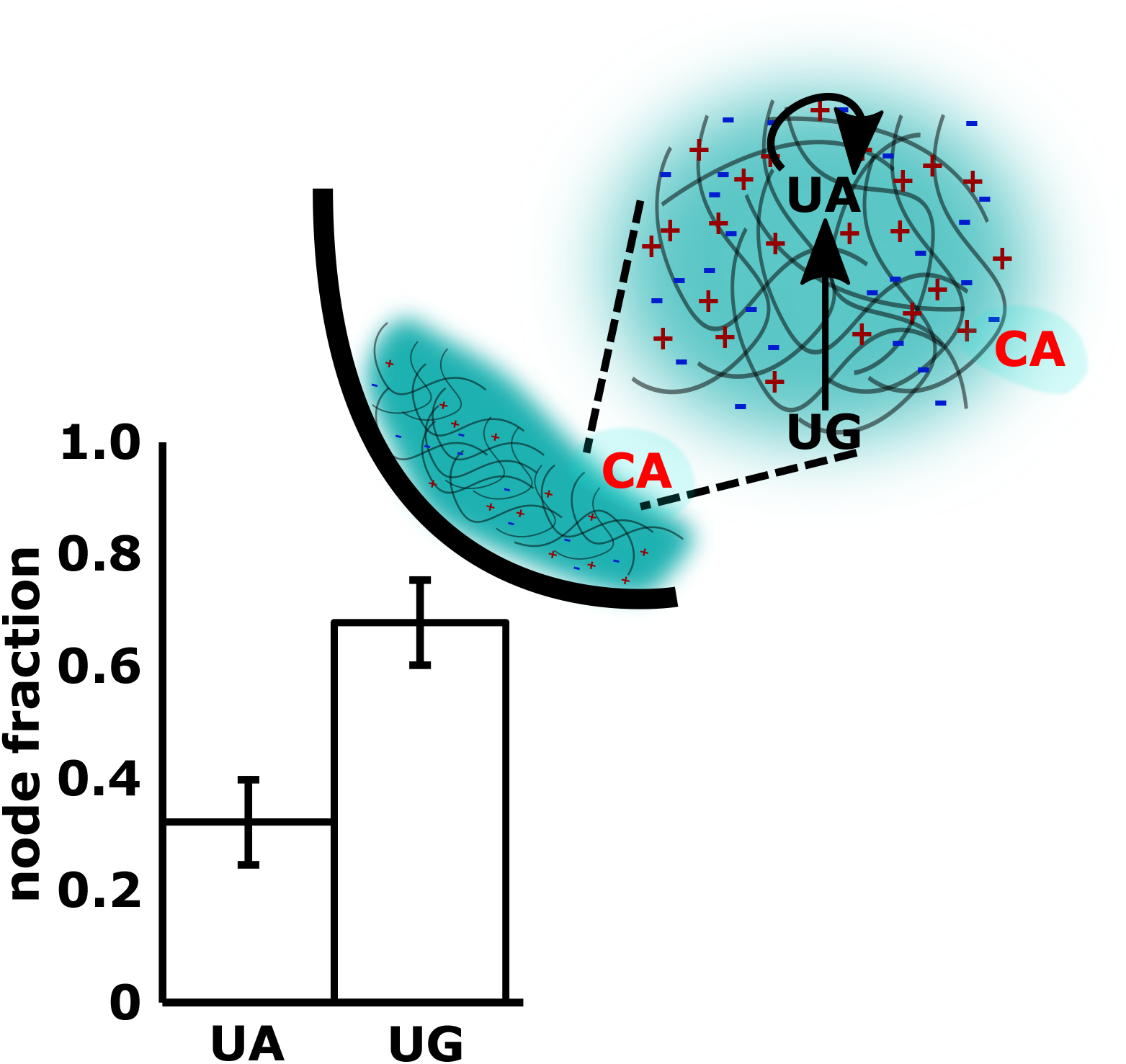
Transient compositional robustness of cross-catalytic network in the coacervate macrophase. The composition of a two node network is subjected to perturbation by adding a strong catalyzing node (’CA’, due to ‘C’ to ‘G’ link, see Ameta et al. (2021)) after encapsulating the network inside the coacervate macrophase. This is a similar experiment as shown in Fig. 4 but here coacervate samples are incubated for 4 *h* at 48°C. Due to longer incubation timings, here the composition is perturbed to the same extent as shown in Fig. 4A. All the measurements are done in triplicates and mean WXYZ product formation is plotted as fraction w.r.t. each node along with standard deviation.

## Bibliography

Adamski, P., Eleveld, M., Sood, A., Kun, Á., Szilágyi, A., Czárán, T., Szathmáry, E., and Otto, S. From self-replication to replicator systems en route to de novo life. Nat. Rev. Chem., 4 (8):386–403, 2020.

Ameta, S., Arsène, S., Foulon, S., Saudemont, B., Clifton, B. E., Griffiths, A. D., and Nghe, P. Darwinian properties and their trade-offs in autocatalytic RNA reaction networks. Nat. Commun., 12(1), 2021.

Ameta, S., Matsubara, Y. J., Chakraborty, N., Krishna, S., and Thutupalli, S. Self-Reproduction and Darwinian Evolution in Autocatalytic Chemical Reaction Systems. Life, 11(4):308, 2021a.

Arsène, S., Ameta, S., Lehman, N., Griffiths, A. D., and Nghe, P. Coupled catabolism and anabolism in autocatalytic rna sets. Nucleic Acids Res., 46(18):9660–9666, 2018.

Axelrod, D., Koppel, D., Schlessinger, J., Elson, E., and Webb, W. W. Mobility measurement by analysis of fluorescence photobleaching recovery kinetics. Biophysical Journal, 16(9): 1055–1069, 1976.

Cakmak, F., Grigas, A. T., and Keating, C. D. Lipid Vesicle-Coated Complex Coacervates. Langmuir, 35:7830–7840, 2019.

Chen, I. A. and Walde, P. From self-assembled vesicles to protocells. Cold Spring Harb. Perspect. Biol., page a002170, 2010.

Crick, F. H. The origin of the genetic code. J. Mol. Biol., 38(3):367–379, dec 1968.

Deamer, D. W. and Georgiou, C. D. Hydrothermal Conditions and the Origin of Cellular Life. Astrobiology, 15:1091–1095, 2015.

Drobot, B., Iglesias-Artola, J. M., Le Vay, K., Mayr, V., Kar, M., Kreysing, M., Mutschler, H., and Tang, T. D. Compartmentalised rna catalysis in membrane-free coacervate protocells. Nat. Commun., 9(1):1–9, 2018.

Dyson, F. Origins of Life. Cambridge University Press, 1985.

Eigen, M. and Schuster, P. A principle of natural self-organization - Part A: Emergence of the hypercycle. Naturwissenschaften, 64(11):541–565, nov 1977.

Eigen, M. Selforganization of matter and the evolution of biological macromolecules. Die Naturwissenschaften, 58(10):465–523, 1971.

Gilbert, W. Origin of life: The RNA world. Nature, 319:618, 1986.

Godfrey-Smith, P. Conditions for evolution by natural selection. J. Philos., 104(10):489–516, 2007.

Haldane, J. The origin of life. Ration. Annu., 148:3–10, 1929.

Hayden, E. J. and Lehman, N. Self-Assembly of a Group I Intron from Inactive Oligonucleotide Fragments. Chem. Biol., 13(8):909–918, aug 2006.

Hayden, E. J., von Kiedrowski, G., and Lehman, N. Systems chemistry on ribozyme self-construction: evidence for anabolic autocatalysis in a recombination network. Angew. Chem. Int. Ed., 120(44):8552–8556, 2008.

Hazen, R. M. and Sverjensky, D. A. Mineral Surfaces, Geochemical Complexities, and the Origins of Life. Cold Spring Harb. Perspect. Biol., page a002162, 2010.

Hordijk, W. and Steel, M. Autocatalytic sets extended: Dynamics, inhibition, and a generalization. J. Sys. Chem., 3(1):1–12, aug 2012.

Hordijk, W., Steel, M., and Kauffman, S. The Structure of Autocatalytic Sets: Evolvability, Enablement, and Emergence. Acta Biotheoretica, 60(4):379–392, dec 2012.

Ianeselli, A., Tetiker, D., Stein, J., Kühnlein, A., Mast, C. B., Braun, D., and Tang, T.-Y. D. Non-equilibrium conditions inside rock pores drive fission, maintenance and selection of coacervate protocells. Nat. Chem., 14:32–39, 2022.

Iglesias-Artola, J. M., Drobot, B., Kar, M., Fritsch, A. W., Mutschler, H., Tang, D. T.-Y., and Kreysing, M. Nat. Chem., 14:407–416, 2022.

Jayathilaka, T. S. and Lehman, N. Spontaneous Covalent Self-Assembly of the *Azoarcus* Ribozyme from Five Fragments. ChemBioChem, 19(3):217–220, feb 2018.

Jia, T. Z., Hentrich, C., and Szostak, J. W. Rapid RNA Exchange in Aqueous Two-Phase System and Coacervate Droplets. Orig. Life Evol. Biosph., 44(1):1–12, feb 2014.

Joyce, G. F. and Szostak, J. W. Protocells and RNA self-replication. Cold Spring Harb. Perspect. Biol., 10(9), sep 2018.

Kauffman, S. A. Autocatalytic sets of proteins. J. Theor. Biol., 119(1):1–24, mar 1986.

Krieger, M. S., Sinai, S., and Nowak, M. A. Turbulent coherent structures and early life below the kolmogorov scale. Nature communications, 11(1):1–14, 2020.

Kumar, M., Singh, A., Secco, B. D., Baranov, M. V., Bogaart, G. v. d., Sacanna, S., and Thutupalli, S. Assembling anisotropic colloids using curvature-mediated lipid sorting. Soft Matter, 2022.

Le Vay, K., Song, E. Y., Ghosh, B., Tang, T.-Y. D., and Mutschler, H. Enhanced ribozyme-catalyzed recombination and oligonucleotide assembly in peptide-rna condensates. Angewandte Chemie International Edition, 60(50):26096–26104, 2021.

Lewontin, R. C. The units of selection. Annu. Rev. Ecol. Evol. Syst., pages 1–18, 1970.

Martin, W., Baross, J., Kelley, D., and Russell, M. J. Hydrothermal vents and the origin of life. Mat. Rev. Microbiol., 6:805–814, 2008.

Martin, N. ChemBioChem, 20(20):2553–2568, oct 2019.

Matsumura, S., Kun, Á., Ryckelynck, M., Coldren, F., Szilágyi, A., Jossinet, F., Rick, C., Nghe, P., Szathmáry, E., and Griffiths, A. D. Science, 354(6317):1293–1296, dec 2016.

Menor-Salván, C. and Marín-Yaseli, M. R. Prebiotic chemistry in eutectic solutions at the water-ice matrix. Chem. Soc. Rev., 41:5404–5415, 2012.

Nakashima, K. K., Vibhute, M. A., and Spruijt, E. Front. Mol. Biosci., 6(APR), 2019.

Nakashima, K. K., van Haren, M. H., Andre, A. A., Robu, I., and Spruijt, E. Active coacervate droplets are protocells that grow and resist ostwald ripening. Nature Communications, 12(1):3819, 2021.

Nghe, P., Hordijk, W., Kauffman, S. A., Walker, S. I., Schmidt, F. J., Kemble, H., Yeates, J. A., and Lehman, N. Prebiotic network evolution: Six key parameters. Mol. Biosyst., 11(12): 3206–3217, nov 2015.

Oparin, A., Braunshtein, A., and Pasynkii, A. The Origin of Life on the Earth. Academic Press, 1957. ISBN 9781483222400.

Poudyal, R. R., Guth-Metzler, R. M., Veenis, A. J., Frankel, E. A., Keating, C. D., and Bevilac-qua, P. C. Template-directed rna polymerization and enhanced ribozyme catalysis inside membraneless compartments formed by coacervates. Nat. Commun., 10(1):1–13, 2019.

Powner, M. W., Gerland, B., and Sutherland, J. D. Synthesis of activated pyrimidine ribonu-cleotides in prebiotically plausible conditions. Nature, 459(7244):239–242, 2009.

Press, W. H., Teukolsky, S. A., Vetterling, W. T., and Flannery, B. P. Numerical recipes 3rd edition: The art of scientific computing. Cambridge university press, 2007.

Reinhold-Hurek, B. and Shub, D. A. Self-splicing introns in tRNA genes of widely divergent bacteria. Nature, 357(6374):173–176, 1992.

Salditt, A., Karr, L., Salibi, E., Le Vay, K., Braun, D., and Mutschler, H. Preprint. doi: 10.21203/rs.3.rs-1989787/v1.

Salibi, E., Peter, B., Schwille, P., and Mutschler, H. Preprint. doi: 10.21203/rs.3.rs-2014540/v1.

Schindelin, J., Arganda-Carreras, I., Frise, E., Kaynig, V., Longair, M., Pietzsch, T., Preibisch, S., Rueden, C., Saalfeld, S., Schmid, B., Tinevez, J. Y., White, D. J., Hartenstein, V., Eliceiri, K., Tomancak, P., and Cardona, A. Fiji: An open-source platform for biological-image analysis. Nat. Methods, 9(7):676–682, jul 2012.

Segré, D., Ben-Eli, D., and Lancet, D. Compositional genomes: prebiotic information transfer in mutually catalytic noncovalent assemblies. Proc. Natl. Acad. Sci. U.S.A., 97(8):4112–4117, 2000.

Strulson, C. A., Molden, R. C., Keating, C. D., and Bevilacqua, P. C. RNA catalysis through compartmentalization. Nat. Chem., 4(11):941–946, nov 2012.

Szathmáry, E. and Demeter, L. Group selection of early replicators and the origin of life. J. Theor. Biol., 128(4):463–486, oct 1987.

Szer, W. Effect of di- and polyamines on the thermal transition of synthetic polyribonu-cleotides. Biochem. Biophys. Res. Commun., 22(5):559–564, mar 1966.

Vaidya, N., Manapat, M. L., Chen, I. A., Xulvi-Brunet, R., Hayden, E. J., and Lehman, N. Spontaneous network formation among cooperative RNA replicators. Nature, 491(7422): 72–77, nov 2012.

van Harren, M., Nakashima, K., and Spruijt, E. Coacervate-based protocells: Integration of life-like properties in a droplet. J. Sys. Chem., 8:107–120, 2020.

Vasas, V., Fernando, C., Santos, M., Kauffman, S., and Szathmáry, E. Evolution before genes. Biology Direct, 7(1):1, jan 2012.

von Kiedrowski, G. Minimal Replicator Theory I: Parabolic Versus Exponential Growth. Bioorganic Chemistry Frontiers, pages 113–146, 1993.

Woese, C. R. The genetic code :the molecular basis for genetic expression. Proc. Natl. Acad. Sci. U.S.A, 1967.

Wołos, A., Roszak, R., Żądło Dobrowolska, A., Beker, W., Mikulak-Klucznik, G., B.and Spól-nik, Dygas, M., Szymkuć, S., and Grzybowski, B. Synthetic connectivity, emergence, and self-regeneration in the network of prebiotic chemistry. Science, 369:eaaw1955, 2020.

Yeates, J. A., Hilbe, C., Zwick, M., Nowak, M. A., and Lehman, N. Dynamics of prebiotic RNA reproduction illuminated by chemical game theory. Proc. Natl. Acad. Sci. U.S.A, (18): 5030–5035, may 2016.

Yeates, J. A., Nghe, P., and Lehman, N. Topological and thermodynamic factors that influence the evolution of small networks of catalytic RNA species. RNA, 23(7):1088–1096, jul 2017.

Zwicker, D., Seyboldt, R., Weber, C. A., Hyman, A. A., and Jülicher, F. Growth and division of active droplets provides a model for protocells. Nat. Phys., 13(4):408–413, apr 2017.

## Bibliography

Kamimura, A., Matsubara, Y. J., Kaneko, K., and Takeuchi, N. PLoS Comput. Biol., 15:1–15, 2019.

